# YAP/TAZ as a Novel Regulator of cell volume

**DOI:** 10.1101/528133

**Authors:** Nicolas A. Perez-Gonzalez, Nash D. Rochman, Kai Yao, Jiaxiang Tao, Minh-Tam Tran Le, Shannon Flanary, Lucia Sablich, Ben Toler, Eliana Crentsil, Felipe Takaesu, Bram Lambrus, Jessie Huang, Vivian Fu, Andrew J. Holland, Steven An, Denis Wirtz, Kun-Liang Guan, Sean X. Sun

## Abstract

How mammalian cells regulate their physical size is currently poorly understood, in part due to the difficulty of accurately quantifying cell volume in a high throughput manner. Here, using the fluorescence exclusion method, we demonstrate that the mechanosensitive transcriptional regulators YAP (Yes-associated protein) and TAZ (transcriptional coactivator with PDZ-binding motif) are novel regulators of single cell volume. We report that the role of YAP/TAZ in cell volume regulation must go beyond its influence on total cell cycle duration or the cell shape to explain the observed changes in volume. Moreover, for our experimental conditions, volume regulation by YAP/TAZ is independent of mTOR. Instead, we find YAP/TAZ directly impacts the cell division volume. Based on the idea that YAP/TAZ is a mechanosensor, we find that inhibiting the assembly of myosin and cell tension slows cell cycle progression from G1 to S. These results suggest that YAP/TAZ and the Hippo pathway may be modulating cell volume in combination with cytoskeletal tension during cell cycle progression.

## Introduction

The question of cell size is at the core of how organisms coordinate cell growth and proliferation. Cell volume dysregulation has been broadly used as a biophysical marker for disease, notably cancer (Kozma and Thomas, 2002; Dannhauser et al., 2017). At a basic level, with increasing cell size, the cell surface to volume ratio shrinks, potentially altering the ratio of membrane-bound components to cytoplasmic components, thus fundamentally changing both inter and intra-cellular dynamics of the cell. Recently, substantial progress has been made towards understanding cell volume regulation, largely enabled by the development of quantitative tools to directly monitor cell cycle progression (Sakaue-Sawano et al., 2008), cell dry mass (Mir et al., 2010; Sung et al., 2013), buoyant cell mass (Son et al., 2012), cell total protein content (Kafri et al., 2013; Ginzberg et al., 2018) and single cell volume in normal culture (Varsano et al., 2017; Cadart et al., 2018; Guo et al., 2017, Wang et al. 2018). Here, to study cell volume in a high throughput manner, we use the fluorescence exclusion method developed by Bottier et al. and others (Cadart et al., 2017; Bottier et al., 2011; Perez Gonzalez, et al., 2018). Using this robust and accurate method, it was revealed that mitotic cells swell suddenly before cytokinesis (Zlotek-Zlotkiewict et al., 2015), and that some types of cells show an adder-like behavior to achieve cell volume homeostasis (Cadart et al., 2018).

In previous work, we demonstrated a relationship between cell volume, cell cortical tension (measured by phosphorylated myosin light chain (pMLC)), and YAP/TAZ activity (measured by nuclear portion of YAP/YAZ) (Perez Gonzalez, et al., 2018; Wang et al., 2018). YAP/TAZ has been previously reported as a key regulator in organ size control (Zhao et al., 2007; Zhao et al., 2008; Tumaneng et al., 2012) and in mechanotransduction (Dupont et al., 2011; Chang et al., 2018; Piccolo et al., 2014). We reported that the mean YAP/TAZ activity is a good predictor of mean cell volume across different cell lines and substrates of varying stiffness. Here, we demonstrate using single cell volume measurements that YAP/TAZ plays an important role in cell volume regulation. The relationship between the Hippo pathway and morphological changes of the cell has already been hinted at in the literature, largely relying on flow cytometry and visual inspection (Horie et al., 2016; Pouffle et al., 2018). In this work, we measure cell volume of human embryonic kidney cells across a group of Hippo pathway CRISPR knockout cell lines with varying degrees of YAP/TAZ activity and demonstrate that the cell volume is positively correlated with YAP/TAZ activity. Moreover, the role of YAP/TAZ in cell volume regulation must not be limited to its influence on the total cell cycle time or the cell shape, nor through its connection with the mTOR pathway. We show that YAP/TAZ directly impacts cell division volume. Since YAP/TAZ is a central player in mechanotransduction (Dupont et al., 2011; Chang et al., 2018; Piccolo et al., 2014), we found that YAP/TAZ activity is directly correlated with cell cytoskeletal tension, and modulating cell tension can delay cell cycle progression. These results suggest that cell tension and the Hippo pathway work together to control the G1/S cell cycle checkpoint, thus determining the cell volume.

## Results

We used the fluorescence exclusion method to quantify single cell volume as previously described (Perez Gonzalez, Rochman, Tao et al., 2018). Briefly, we fabricated microchannels coated with collagen I (Fig. 1a). Single cells were seeded at low density and allowed to adhere. After cell adhesion, medium was infused with fluorescent FITC-Dextran, which evenly labeled the cell surroundings but remained exclusively in the extracellular medium, exhibiting minimal endocytosis within 5 hours (Fig. 1b). The epifluorescent images obtained were then segmented (Fig. 1c), and the 3D cell volume was computed as depicted in Fig. 1d. and reconstructed (Fig. S1a-c)

**Figure 1.**
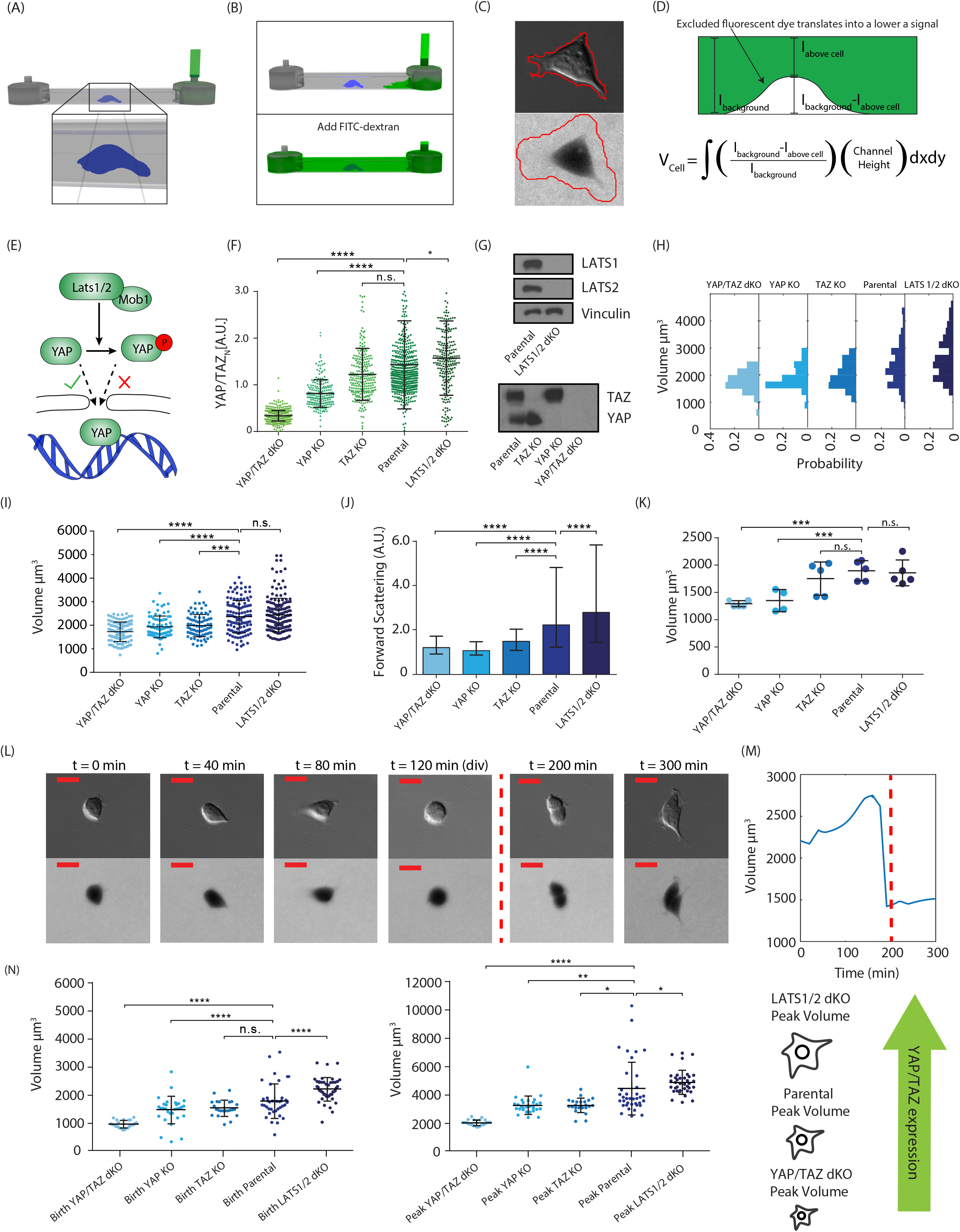
(a-d) Cartoons depicting Fluorescence Exclusion (FX) method. (a) Side view of microdevice. (b) Cells are seeded in the device before adding the fluorescent dye (top). The dye is not membrane permeable and therefore is excluded from the cell interior (bottom). (c) Sample DIC and Fluorescent image for volume calculation. (d) Conversion of fluorescent image into volume. (e) Hippo Pathway cartoon, showing key elements of the pathway. (f) Nuclear YAP/TAZ content by quantitative Immunofluorescence. Hippo pathway activity was assessed by qIF in the parental line and all CRISPR Knockouts (N_YAPTAZ dKO_=223, N_YAP KO_=156, N_TAZ KO_=287, N_HEK 293A_=331, N_LATS1/2 dKO_=207). (g) Hippo pathway Knockout validation. Western Blots were performed to assess YAP and TAZ expression in all CRISPR Knockouts. (h) Volume measurements of Hippo Pathway Knockouts via FX method. Panel shows the distribution for each cell line (N_YAPTAZ dKO_=130, N_YAP KO_=118, N_TAZ KO_=162, N_HEK 293A_=135, N_LATS1/2 dKO_=185). (i) Volume averages from the FX method. Hippo pathway volume averages from figure (g). (j) Volume measurement of Hippo pathway knockouts via Flow Cytometry. (k) Volume measurement of Hippo pathway knockouts via Coulter Counter. (N_YAPTAZ dKO_=5, N_YAP KO_=4, N_TAZ KO_=5, N_HEK 293A_=5, N_LATS1/2 dKO_=5). (l) Cell volume as a function of time. Cells volume was monitored over time for 5 hours. (m) Volume trajectory. Sample cell growth trajectory is shown as a function of time for a single cell. (n) Daughter cell volume and volume before division. Averages are presented for the parental cell line and all Hippo pathway KOs.

### YAP/TAZ and the Hippo Pathway Regulate Single Cell Volume

Previous work reported positive correlations between the 2D cell adhesion area with cell YAP/TAZ expression (Plouffe et al., 2018), and the 3D cell volume with cell YAP/TAZ activity (defined as nuclear YAP/TAZ content) (Perez Gonzalez, Rochman, Tao et al., 2018). We sought to further elucidate the relationship between the Hippo pathway and cell volume through the utilization of CRISPR knockouts for Hippo genes (Fig. 1e). Other studies have shown that activation of the Hippo pathway is followed by the phosphorylation of YAP/TAZ, leading to cytoplasmic localization and inhibition of YAP/TAZ (Zhao et al., 2007) (Fig. 1e). In this pathway, LATS1/2 is responsible for YAP/TAZ phosphorylation and inactivation, therefore their absence increases YAP/TAZ nuclear localization and subsequent activity. Validations of these CRISPR knockouts were performed via quantitative immunofluorescence (qIF) (Fig. 1f) and Western Blots (Fig 1g). More details on qIF can be found in the SM (Fig. S1d-f). We found that the average nuclear YAP/TAZ activity is increased in the LATS1/2 double knockout (dKO) when compared to the parental HEK 293A, although there is some overlap between the populations. Similarly, the TAZ KO and YAP KO exhibit a significant decrease in nuclear YAP/TAZ. Finally, qIF for the YAP/TAZ dKO showed low YAP/TAZ activity and low variability, suggesting this corresponds to non-specific binding and is an indication of the level of background noise in our measurement.

Across the four CRISPR generated cell lines and the parental HEK 293A, we found that the average amount of nuclear YAP/TAZ strongly correlated with the average cell volume (Fig. 1h,i), as seen before for cells growing on substrates of varying stiffness (Perez Gonzalez, Rochman, Tao et al., 2018). Accordingly, when comparing volume distributions of all 5 cell lines (Fig. 1h), we found that YAP/TAZ activity correlates with an increasing abundance of larger cells, e.g., LATS1/2 dKO has mostly larger cells while YAP/TAZ dKO has mostly smaller cells. When comparing the average cell volume between populations, we observed an increase of 3.01 % in the LATS1/2 dKO. As YAP/TAZ expression decreased, we observed a volume decrease of 15.8% in the TAZ KO, 18.2% in the YAP KO and 27.2% in the YAP/TAZ dKO (Fig. 1i). It is worth noting that under favorable culture conditions, LATS1/2 are largely inhibited and YAP/TAZ are active. This may explain why LATS1/2 dKO only produced a modest effect on cell size whereas YAP/TAZ KO produced more dramatic effects. Additionally, we assessed the cell spreading area, noticing that volume and area are positively correlated with each other, but in some cases, a volume change could be observed in the absence of an area change (Fig. S1g,h). We also used two more common approaches to assess cell size: Coulter counter (Fig. 1k) (Burke et al., 2012; Conlon et al., 2001) and flow cytometry (Fig. 1j) (Plouffe et al., 2018). Despite requiring the cells to be resuspended, grossly changing their morphology, both the Coulter counter measurements (Fig. 1j) and flow cytometry measurements showed a similar cell volume trend (Fig. 1k).

In addition to the static cell volume, we examined how the cell volumetric growth rate as well as the volume at the beginning and end of the cell cycle varies with the presence of YAP/TAZ. We tracked single cell volume for five hours within the fabricated microchannels described above (Fig. 1l) and obtained cell growth trajectories (Fig. 1m) for each of the five cell lines. A portion of cells undergoing mitosis were observed, for which we measured the volume immediately before and after division (referred to as division and birth volumes onwards). We found that both the cell birth volume and the division volume increased with increased activity of YAP/TAZ (Fig 1n), explicitly confirming that the observed volumetric change corresponds to an intrinsic change in volume within the population.

### Single cell growth rate is proportional to cell size and follows a universal growth law

To further understand single cell growth, we quantified added volume per unit time (volumetric growth rate) for many single cells. Fig. 2a shows the average cell growth trajectory in blue and the standard deviation as the gray area for each of the 5 cell lines. Fig. 2a also displays sample trajectories for each of the cell lines as an inset on the top right of each figure. The degree to which these cell lines display obvious mitotic swelling as previously reported in the literature (Zlotek-Zlotkiewicz et al., 2015) varies with YAP/TAZ activity. The TAZ KO, parental cell line, and LATS1/2 dKO display a sharp volume increase and decrease right before division. However, the YAP KO and YAP/TAZ dKO show reduced or unresolvable mitotic swelling at the time resolution used in this study, suggesting that YAP may play a role in mitotic swelling. To quantify growth trajectories and obtain the growth rate, we fitted an exponential growth law to each trajectory, obtaining *dV/dt* versus *V* for each cell (Fig. 2b, 2c). Additionally, we fitted linear growth curves to the same data (Fig. S2a,b) since it has been noted that it is difficult to distinguish between an exponential growth law from a linear growth law (Ginzberg et al., 2015) largely due to the small range of volumes observed within a single cell cycle (roughly a two-fold variation). Indeed, in the next section, we discuss that the growth law is unlikely to have a measurable influence on the average cell volume; however, our results show that the volumetric growth rate is proportional to the cell volume across populations. When all growth trajectories for all cells are overlaid (Fig. 2d), it is apparent that all 5 cell lines follow a similar growth law, i.e., the growth rate *dV/dt* ∝*V* regardless of YAP/TAZ activity (and regardless of whether *dV/dt* for each cell was fitted assuming an exponential or linear dependence on *V*). The combination of multiple cell lines spanning a much larger range of volumes lends greater confidence to the observation that across populations there is a strong linear dependence of *dV/dt* on *V,* indicating an exponential volumetric growth rate with respect to cell volume.

**Figure 2.**
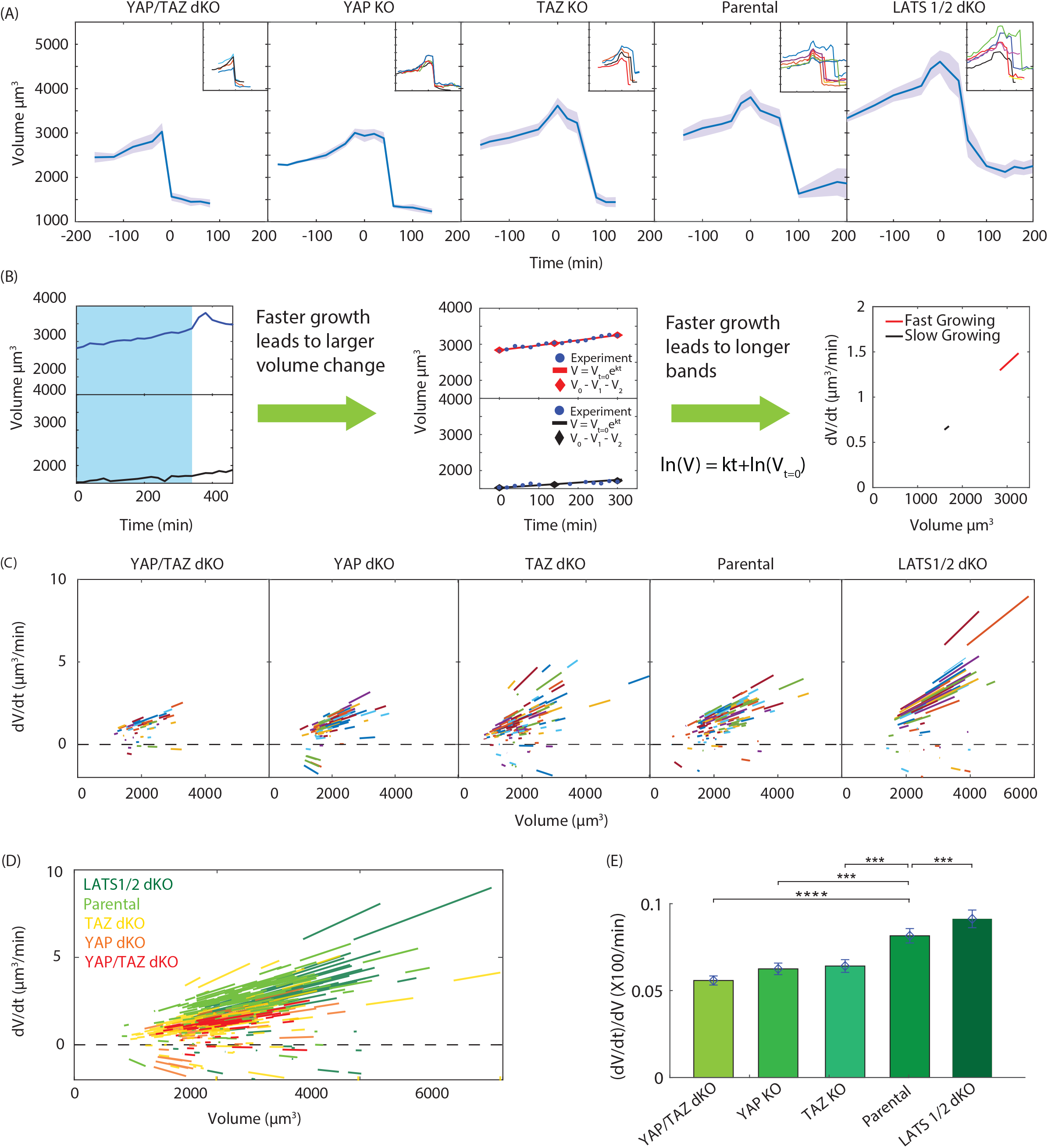
Average cell growth trajectory. The average trajectory is shown in blue and the first standard deviation is shown by the gray area. Sample cell growth trajectories are displayed in an inset to qualitatively show their behavior (b) Extraction of growth rate, dV/dt, as a function of time. Left panel shows the sample trajectories of two cells, a small cell (bottom panel) and a big cell (top panel). For each cell, we use three points in order to determine the constant for exponential growth (middle panel). Sample plot of single cell growth rate versus the volume of that cell (right panel). (c) Growth rate versus cell at single cell level. (N_YAPTAZ dKO_=130, N_YAP KO_=118, N_TAZ KO_=162, N_HEK 293A_=175, N_LATS1/2 dKO_=185). (d) Growth rate curves overlapped for all Hippo knockouts. With increasing YAP/TAZ activity, the growth rate is slightly higher for the same cell volume. (e) The average slope of *dV/dt* versus V.

The data shows that the presence of YAP/TAZ seems to be regulating *dV/dt* with increasing YAP/TAZ activity associated with slightly higher *dV/dt*. At first glance, this might seem to be the critical observation needed to explain how YAP/TAZ and Hippo are regulating cell volume. Thus, it is natural to focus on the regulation of *dV/dt*. However, average cell volume does not depend only on growth rate but also the duration of the cell cycle, *τ*. It can be shown theoretically that the average cell volume is slightly less than *3/2τdV/dt* (see SM for details) where *τ* is the average cell cycle duration. YAP/TAZ has also been shown to regulate the duration of the cell cycle (Pouffle, 2018) (see Fig. 3a) with increasing YAP/TAZ activity associated with smaller *τ*. It is clear that YAP/TAZ is affecting *dV/dt* and *τ* in opposite ways (and examining the trend in *τ* alone is also insufficient to explain volume regulation – see the following section). A larger increase is observed for *dV/dt* than what would be necessary to compensate for the observed decrease in *τ* and keep the average volume constant. How YAP/TAZ skews this balance in favor of increased *dV/dt* remains unclear, and more information regarding the role of YAP/TAZ in cell cycle regulation may be needed to understand this phenomenon (see Fig. 5 for more discussion).

**Figure 3.**
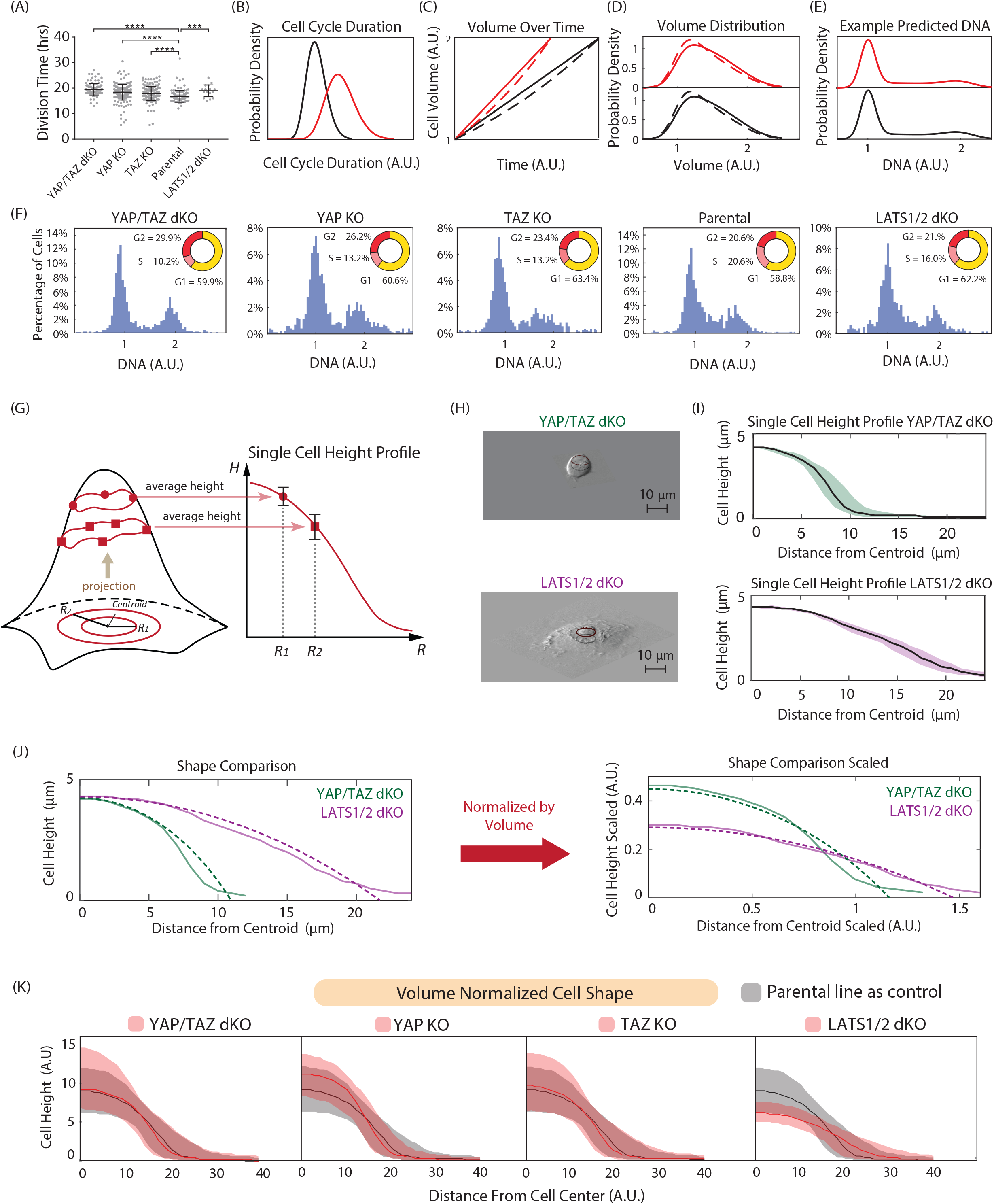
Volume difference across Hippo pathway knockouts is independent of cell growth dynamics and geometric changes. (a) YAP/TAZ Decreases Cell Cycle Duration and Doubling Time. Cell cycle duration and bulk doubling time for the HEK 293A and five KO lines. (N_YAPTAZ dKO_=107, N_YAP KO_=216, N_TAZ KO_=186, N_HEK 293A_ =122, N_LATS1/2 dKO_=18). (b) Two populations with different cell cycle duration are considered using a mathematical model (SM). Long cell cycle duration and short cell cycle duration. (c) Two growth schemes are considered, linear or exponential growth. A solid red line represents linear growth with short cell cycle duration and a dashed red line represents exponential growth with short cell cycle duration. The solid black line represents linear growth with long cell cycle and a dashed black line represents exponential growth with long cell cycle duration. (d) Volume distributions for each case. (e) Computed DNA distribution. Predicted from the mathematical model in SM. (f) YAP/TAZ Does Not Change the Cell Cycle Phase Distribution. A comparison of the DNA distributions from Hoestch staining across the cell lines showed no significant difference (inset cell cycle fraction). (N_YAPTAZ dKO_=1137, N_YAP KO_=961, N_TAZ KO_=774, N_HEK 293A_=1202, N_LATS1/2 dKO_=876). (g) The Role of YAP/TAZ in 3D Cell Shape – cartoon of analysis. Cartoon depicts single cell profile from volume data (h) 3D rendering. Two sample cells were considered to demonstrate the three-dimensional rendering. Top panel shows the YAP/TAZ dKO and the bottom panel shows the LATS1/2 dKO. (i) Sample profile for two cell lines. Top panel shows YAP/TAZ dKO and bottom panel shows LATS1/2 dKO. Shaded area is median 50%. (j) Cell shape comparison. Left panel shows the cell height distribution as a function of the radial distance from the cell’s center of mass (volume in this case). Right panel shows the comparison when scaled by volume. The original and volume normalized apical cell shapes with the median height function (solid lines) and best fit spherical cap (dashed lines) see SM for details on cap fitting. (k) Volume Difference Is Not Explained by Cell Shape. Top panel shows volume normalized median cell height distributions across each ensemble with the indicated KO in red and HEK293 in gray. Shaded area is median 50%. Only the LATS1/2 dKO shows a significant difference in 3D apical shape.

Due to endocytosis of FITC-Dextran on longer time scales, we are not able to obtain growth trajectories for the complete cell cycle. Therefore, we are unable to observe correlations between the birth volume and the added volume at the single cell level; however, we do observe a population of cells that exhibit near zero growth over 5 hours, which we interpret as quiescent cells. We observed that cells in quiescence can also transition out of quiescence and grow again. Finally, it is worth noting that growing cells show a continuous and proportional growth rate vs. volume curve with no visible dependence on cell cycle phase.

### Volume differences across Hippo pathway knockouts are not explained by cell cycle duration or volumetric growth law

It is clear how changing the birth and division volumes may affect the average population volume; however, it is also possible that the cell cycle duration, cell cycle phase distribution, and growth rate as a function of volume could impact the mean volume while keeping the birth and division volumes constant. For example, suppose we have a hypothetical cell for which *dV/dt* is positive during G1 and zero during S and G2. Further suppose that we can lengthen the duration of G2 while keeping the birth and division volumes constant. For that cell, lengthening G2 will increase the average volume of the population since each cell will spend more time at its maximum volume. We did not observe growth laws of this type in the cell lines utilized for this paper; however, we sought to examine how small variations in the growth law (e.g., comparing the exponential and linear models) may affect average volume. In previous work (Rochman, Popescu and Sun, 2018), we discussed how some conserved quantities may be used to calculate population averages of age-dependent measurements. In particular, volume and DNA distributions for an ensemble may be calculated given the growth trajectory, V(t), and DNA content progression, DNA(t), (see SM for details). We may compare two cell volume distributions: one obtained with a long cell cycle duration (Fig. 3b red curve) and one obtained with a short duration (Fig. 3b black curves). The V(t) curve for the ensemble with the shorter duration will have a steeper rise regardless of whether the growth law is linear or exponential (Fig. 3c); however, the resultant volume distributions will be negligibly different (and the difference depends only on whether the growth rate is linear or exponential, not on the cell cycle duration) on the scale of variation we observe across the CRISPR knockouts (Fig. 3d), see SM for details. When the ratio of time spent in each cell cycle phase G1, S and G2 is conserved, as observed across the CRISPR knockouts (Fig. 3f), we also predict the DNA distribution to be independent of cell cycle duration (Fig. 3e). Thus, cell cycle duration is not predicted to impact the volume distribution, and while modifying the growth law may modestly change the mean cell cycle duration, only extremely nonlinear trends could replicate the magnitude of variation we observe across the CRISPR knockouts. As we see from Fig. 2d, the growth laws for all cell lines are generally similar, and the cell cycle duration for 5 cell lines are also very similar (Fig. 3a). Therefore, any impact the Hippo pathway and YAP/TAZ activity may have on the volumetric growth law or cell cycle duration (Fig. 3a) alone is unable to explain the volume variation across the CRISPR knockouts.

### Volume variations are not explained by cell geometry

In a previous paper (Perez Gonzalez, Rochman, Tao et al., 2018), we have shown that cell geometry plays a major role in cortical force balance and is an important factor in determining cell size. In principle, regulating the spatial distribution of cortical contractility while conserving the osmotic gradient across the cell cortex could result in large changes in cell volume. For example, suppose increasing YAP/TAZ activity decreases cell protrusivity, resulting in a more hemispherical cell shape and thus higher curvature. This higher curvature, with the same pMLC expression, leads to a larger contractile force. If the pressure gradient across the cortex was constant, one way to balance that larger contractile force would be to increase cell volume and decrease the curvature while maintaining the same shape. Thus, we sought to examine how cell shape varies with YAP/TAZ activity independently of cell volume, and examined the 3D shape where the apical surface was reconstructed from the epifluorescent images used to calculate volume (Fig. 3g, h) and normalized by cell volume (Fig. 3i). We examined both the raw height profiles (Fig. 3j) as a function of the distance from the center of each cell as well as the fitted spherical caps (Fig. S3f). We found that there was no consistent trend in cell shape with increasing YAP/TAZ activity. The YAP/TAZ dKO, YAP KO, TAZ KO, and parental line (HEK293A) were all found to be remarkably self-scaling. Despite having substantially different volumes, these four cell lines all had similar height profiles. It is interesting to note, however, the LATS1/2 dKO does not exhibit the same shape, spreading more than the others (more “pancake-like”). Thus, the role the Hippo pathway and YAP/TAZ activity may play in cell shape regulation alone cannot explain the variations in volume observed across the CRISPR knockouts.

### Cell Volume Regulation by YAP/TAZ is independent of mTOR activity

Another possible explanation for the observed cell volume increase with increasing YAP/TAZ activity is from the potential crosstalk between the Hippo and mTOR pathways. The Hippo pathway has been reported to modulate the mTOR pathway under certain conditions. (Tumaneng et al., 2012) showed that in mouse tissue, overexpression of YAP leads to the upregulation of AKT through tumor suppressor PTEN, and thus activates mTORC1/2 and its downstream components. Over-expression of YAP also increased the level of phosphorylation of both S6K Thr 389 and AKT Ser 473 in tissue. As mTOR has been described as a major regulator of mammalian cell size due to its relationship with amino acid import and protein synthesis (Fingar et al., 2002; Yang et al., 2007; Lloyd, 2013), it is natural to ask whether on the single cell level the regulation of cell volume by the Hippo pathway is mediated by the crosstalk between Hippo and mTOR. If YAP/TAZ activity also modulates mTOR activity on the single cell level, it may explain the observed cell volume variation in Hippo pathway KOs. In order to explore this potential crosstalk between YAP/TAZ and the mTOR pathway, we first inhibited mTOR using rapamycin and replicated the previously published effect of rapamycin on cell size (Fingar et al., 2002) (Fig. S4a-c). After 4hrs of treatment, mTOR activity is diminished as reported by pS6 expression using qIF (Fig 4a, b). However, cell volume did not change significantly after 4 hrs. Instead, a noticeable cell volume decrease of 28.9% occurred after 72 hours, close to the range previously reported (Fingar et al., 2002). In addition, we found that under mTOR inhibition, YAP/TAZ activity remained unchanged, suggesting that mTOR is not an upstream regulator of YAP/TAZ (Fig. S4a,b,d,e).

**Figure 4.**
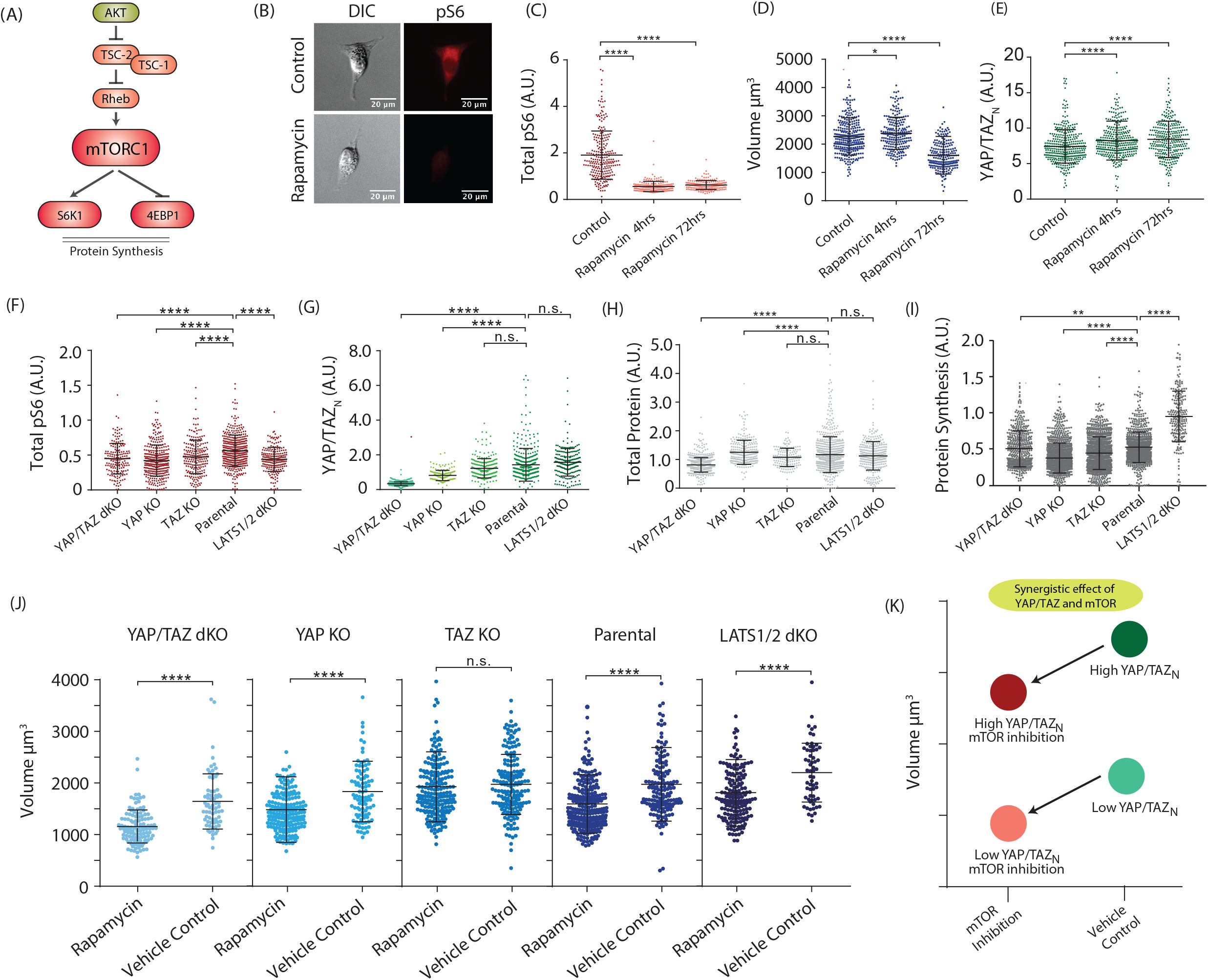
mTOR pathway and cell volume. Description of the key elements of the pathway. (b) Sample images of control cells and rapamycin treatment. Left two show sample HEK293A cells from control experiment whereas two right images show sample cells after rapamycin treatment (c) mTOR inhibition confirmation via qIF. Expression of ribosomal activity reported via qIF of ribosomal protein pS6 (N_Control_=248, N_Rapamycin-4hours_ =196, N_Rapamycin-72hours_=178). (d) Rapamycin treatment decreases cell volume after 72 hours of treatment. Cell volume for control cells and rapamycin treated cells at 4 and 72 hours. (N_Control_=314, N_Rapamycin-4hours_=224, N_Rapamycin-72hours_=221) (e) mTOR inhibition does not affect YAP/TAZ activity. Nuclear YAP/TAZ expression measured via qIF (N_Control_=354, N_Rapamycin-4hours_=326, N_Rapamycin-72hours_=296) (f) mTOR activity, measured as pS6 expression, does not change in Hippo pathway knockouts. (N_YAPTAZ dKO_=177, N_YAP KO_=336, N_TAZ KO_=218, N_HEK 293A_=406, N_LATS1/2 dKO_=202). (h) Total protein content does not change with Hippo pathway knockouts. N_YAPTAZ dKO_=664, N_YAP KO_=269, N_TAZ KO_=150, N_HEK 293A_=772, N_LATS1/2 dKO_=314). (i) Protein synthesis rate, measured using SUnSET, for Hippo pathway knockouts. The synthesis rate is generally unchanged except for a slight increase in LATS1/2 KO. (N_YAPTAZ dKO_=481, N_YAP KO_=1027, N_TAZ KO_=962, N_HEK 293A_=783, N_LATS1/2 dKO_=219) (j) Cell volume comparison for Hippo pathway knockouts before and after mTOR inhibition. (N_YAPTAZ dKO - Control_ =70, N_YAPTAZ dKO - Rapamycin_=115, N_YAP KO - Control_= 95, N_YAP KO - Rapamycin_=193, N_TAZ KO - Control_=176, N_TAZ KO - Rapamycin_=206, N_HEK 293A - Control_=154, N_HEK 293A - Rapamycin_=258, N_LATS1/2 dKO - Control_=61, N_LATS1/2 dKO - Rapamycin_=169) (l) Our results suggest synergistic effect of YAP/TAZ and mTOR in cell volume control. Inhibiting mTOR uniformly decreases cell size in all YAP/TAZ knockouts.

**Figure 5.**
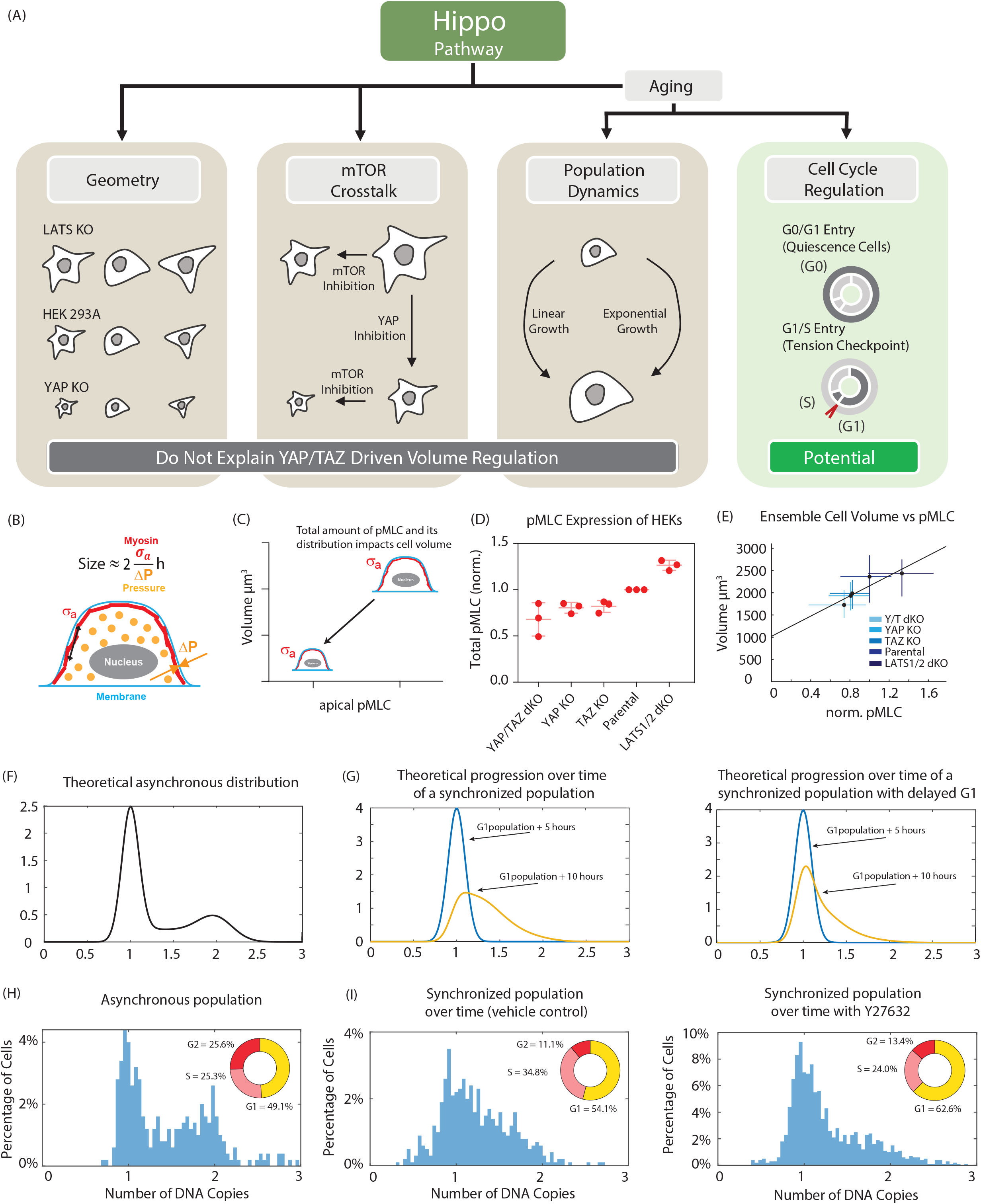
Possible explanations for the mechanism by which the Hippo pathway regulates cell volume. (i) Hippo affects cell geometry: excluded (does not explain cell volume variation). Cells appear to self-scale their shape when YAP and/or TAZ are knocked out. (ii) Hippo cross-talk with mTOR: excluded. In low amino acid conditions this cross-talk is an essential component of both signaling pathways (Tumaneng et al., 2012); however, under the experimental conditions tested for this study, mTOR and Hippo are observed to act independently. (iii) Hippo affects cell volume by influencing cell growth rate and cell cycle distribution: plausible. Potential changes in growth rate caused by perturbing YAP/TAZ fails to explain cell volume regulation; however, cell cycle regulation including modulation of G1/S checkpoint may explain cell volume variation. (b) Cartoon showing relationship between volume and cortical tension, σ_a_. (c) Cartoon showing expected relationship between apical pMLC and volume. (d) qIF measurement of pMLC Expression Changes Among HEKs. Each data point is a biological repeat. Each point is an average of 100-200 cells. (e) Linear relationship between average cell volume and average cell pMLC. (g) Theoretical prediction of DNA distribution over the course of the experiment. See SM. (h) Progression from G1 after shake-off synchronization in M. Left panel shows the synchronized population (blue) versus the population evolving over time (yellow). Right panel shows the synchronized population (blue) and the population evolving over time when G1 progression is delayed (yellow) (i) asynchronous cell cycle distribution for HEK 293A. (j) Y27632 assay. Left panel shows how the cells progress from a synchronized G1 population with vehicle control (DMSO). Right panel shows how the cells progress from a synchronized G1 population with Y27632. Insets show the fraction of cells in each cell cycle phase.

Next, we sought to characterize mTOR activity (reported to be involved in protein production) in all Hippo pathway KOs, as well as the total cell protein content and cell protein synthesis rate. Total cell protein content is measured using the total fluorescence from cells stained with a succinimidyl ester dye (Kafri et al., 2013). This measurement has already been reported to be well correlated with measurements of dry mass by quantitative phase microscopy (Kafri et al., 2013). The cell protein synthesis rate is measured using the SUnSET method (Schmidt et al., 2009). By treating cells with low concentration puromycin and staining puromycin labeled pre-mature peptides, we quantified the rate of protein synthesis in single cells by assessing the qIF signal of puromycin labeled peptides within the cytoplasm. We found that although there is major change in YAP/TAZ expression throughout these cell lines (Fig. 4d), mTOR activity, as measured by total pS6, remained fairly constant (Fig. 4e), further ruling out crosstalk between these pathways at the single cell level. We found no change in total cell protein content across 5 cell lines. The cell protein synthesis rate is also generally constant, with LATS1/2 dKO showing a slight increase. Finally, we asked whether mTOR or Hippo pathways are independent in their regulation of cell volume and whether they have a synergistic effect. We found that when inhibiting mTOR activity with rapamycin in all Hippo pathway KOs, there was a general trend of obtaining additional volume reduction to what we had already seen in the Hippo KOs. Only the TAZ KO showed no significant volume reduction after mTOR inhibition. These results suggest that under the conditions tested, Hippo and mTOR have a synergistic effect in their regulation of cell volume and act essentially independently.

### Tension regulation through the G1/S checkpoint could explain cell volume variations

So far, we have demonstrated that all volume changes observed due to the modulation of YAP/TAZ activity are not explained by the role of YAP/TAZ in the regulation of the volumetric growth rate (Fig. 2), cell cycle duration (Fig. 3), cell geometry (scaled by volume) (Fig. 3), or mTOR activity (Fig. 4). In previous work (Perez Gonzalez, Rochman, Tao et al., 2018), we discussed the relationships between YAP/TAZ, cortical tension, and volume. Here we go a step further to propose that the trend of increasing volume across the CRISPR knockouts might be explained as a consequence of increasing cell tension which in turn increases YAP/TAZ activity (Fig. 5a,b). As previously published, we propose that the distribution of cortical tension, maintained through phosphorylated myosin mediated active contraction, determines the force balance at the cortex, thereby determining the volume of the cell. To validate this framework for the Hippo pathway KOs, we first measured pMLC expression, and found that, as expected, it was positively correlated with YAP/TAZ expression and activity (Fig. 5d) while its distribution in the cell apical cortices remained fairly constant (Fig. S5a-c). When comparing our measured values for pMLC and volume, we found that pMLC was an excellent proxy for cell volume (Fig. 5f). We went on to characterize the pMLC expression for cells identified to be in G1 by Hoechst labelling for each cell line and found that the G1 exit value for pMLC (the value bounding 90% of cells in G1) was again correlated with the average cell volume of the cell line (Fig. S5a). We hypothesize that YAP/TAZ and pMLC may be influencing a G1/S cell cycle checkpoint. To support the existence of such a checkpoint which has recently been motivated in monolayers (Uroz et al., 2018) for isolated HEK single cells, we measured cell cycle progression in a synchronized population under pMLC inhibition with the ROCK inhibitor Y27632. We took an asynchronous population (Fig. 5g), performed mitotic shake off detachment to obtain dividing cells, and let this population progress to G1 (Fig. 5h). We then compared the evolution of this population over time with (Fig. 5j) and without (Fig. 5i) the addition of Y27632. We found that with the addition of Y27632, there were significantly fewer cells progressing from G1 to S. Recently, it was shown that cell cytoskeletal tension impacts the association of the SW1/SNF complex with YAP/YAZ, which inhibits transcriptional activity of nuclear YAP/TAZ (Chang, et al. 2018). This result, in combination with our observation, suggests that pMLC activity and cell tension together with YAP/TAZ activity are involved in the determination of this cell cycle checkpoint.

## Discussion

Our data shows that YAP/TAZ and the Hippo pathway are involved in regulating single cell volume and provides insights into how YAP/TAZ is potentially regulating cell size. We found cell birth volume and division volume to be dependent on YAP/TAZ activity with higher YAP/TAZ activity leading to larger cell size. We validated (Pouffle, 2018) that cell cycle duration decreases with increasing YAP/TAZ activity, and also observed that the volumetric growth rate *dV/dt* increases with YAP/TAZ activity.

Neither the observed dependence on YAP/TAZ activity for the regulation of cell cycle duration nor any potential modulation of the volumetric growth law is able to explain the volume variation observed. We additionally considered the possibility that this phenomenon is purely mechanical, owing to the potential role of YAP/TAZ in the regulation of cell shape; however, we found four out of five of the cell lines that we investigated to exhibit self-similar shapes negating this hypothesis as well. Finally, we sought to establish YAP/TAZ as either directly up or downstream of mTOR, a known regulator of cell volume, and instead, w found them to act both independently and synergistically under the conditions tested in this study.

Having exhausted these possible explanations, we aimed to propose a novel mechanism implicating the role of YAP/TAZ in the regulation of cortical tension and thus cell volume throughout the G1/S checkpoint. We established a strong positive correlation between YAP/TAZ activity, cell volume, and cell tension and went on to validate the importance of cell tension through the navigation of the G1/S checkpoint in isolated HEK single cells by demonstrating that ROCK and myosin inhibition delays or prohibits the exit of G1 and entrance to S. We posit that YAP/TAZ modulates cortical tension, and thus cell volume during this transition, further impacting volume at birth and division. Recently, it was shown that the SWI/SNF complex can inhibit YAP/TAZ transcriptional activity in the cell nucleus in a cell tension sensitive manner (Chang et al., 2018), and YAP/TAZ nuclear import is sensitive to mechanical forces (Elosegui-Artola et al., 2017). Our findings suggest that tension sensitivity of YAP/TAZ ultimately translates to differences in cell cycle progression, providing a novel explanation of cell size control in different mechanical environments.

