## Supplemental Materials for "YAP/TAZ as a Novel Regulator of cell volume"

### YAP/TAZ as Novel Regulators of Cell Volume: Supplementary Material

[1]Department of Chemical and Biomolecular Engineering, Johns Hopkins University, Baltimore, Maryland 21218, USA.[2]Department of Mechanical Engineering, Johns Hopkins University, Baltimore, Maryland 21218, USA. [3]Department of Molecular Biology and Genetics, Johns Hopkins University, School of Medicine, Baltimore, Maryland, 21205, USA. [4]Department of Environmental Health and Engineering, Johns Hopkins Bloomberg School of Public Health, Baltimore, Maryland, 21205, USA. [5]Department of Pharmacology and Moores Cancer Center, University of California, San Diego, La Jolla, California. [6]Physical Sciences in Oncology Center (PSOC), Johns Hopkins University, Baltimore, Maryland 21218, USA

[†]Corresponding author

*Author's contributed equally

**Mathematical Predictions of Cell Volume Distributions**

We identified within the five cell lines observed, that the mean volume of each population increases with increasing YAP/TAZ activity in that population. This is largely due to the fact that the birth and division volumes increase across the populations; however, we also explored ways in which the mean ensemble volume could change while the birth and division volumes remain unchanged. To do so, we utilized the von Foerster equation, which computes the distribution of cells with age-related (cell age is defined as the time that has elapsed since the last cell division) quantities including volume:

$$\frac{\partial n}{\partial t} + \frac{\partial n}{\partial a} = \lambda \qquad (1)$$

Where *n(a,t)da* is the number of cells in the distribution with ages (or time since birth) between *a* and *a+da* and *t* is the chronological time. $\lambda$ represents the loss rate – the rate at which cells leave the ensemble. The number density of cells at age *a+da* at time *t+dt* is simply the number density of cells of age *a* at time *t* less those that left the ensemble due to cell division etc. Considering $\lambda$ to represent only loss due to mitosis we may write the boundary condition:

$$n(0,t) = 2\int_0^\infty \lambda(a)n(a,t)da \qquad (2)$$

which illustrates that the number of cells at age zero is simply twice the number of cells that just divided. We may further rewrite λ in terms of the division time distribution, *w(a),* which is also the cell cycle duration distribution – the amount of time it takes for a cell to divide (Stukalin et al (2013)). The probability of undergoing mitosis per unit time at age *a* is the ratio of the population of cells observed to divide at age *a* over the fraction of cells which have matured to age *a* without yet dividing. Given only *w(a),* one may then solve for the bulk growth rate of the mitotic population, *b*, using the expression (see Stukalin et al (2013) for a derivation):

$$1 = 2\int_0^{\infty} \exp(-ba)w(a)da \qquad (3)$$

Where the cell number distribution is assumed to follow the form $n(a,t) = g(a)\,e^{bt}$, and the age distribution is explicitly:

$$g(a) = 2bexp(-ba)\left[\int_a^{\infty} w(a')da'\right] \qquad (4)$$

In previous work (Rochman et al., 2018) we showed the age distribution to be self-scaling. In other words, the probability of observing a cell of scaled age, $a/\mu$, where $\mu$ is average duration of the cell cycle, is independent of $\mu$. While there is some dependence on the coefficient of variation of the cell cycle duration distribution, given this self-scaling, one may approximate the cell cycle completion distribution, that is the probability that a cell has completed *100*x%* (and where "x" will be denoted the cell "completion-fraction") of its cycle at the time observed as the following conserved function (See Fig. S3b)

$$\rho(x) = 2\ln(2)\exp(-\ln(2)x) \qquad (5)$$

which may be validated through simulation.

This expression may be used to calculate the volume distribution for an ensemble (in principle, any observable e.g. DNA content (Fig. 3e)) given P(V|x), the conditional probability that the volume is some value given that the cycle completion fraction is *x*:

$$\rho(V) = \int_0^1 P(V|x)\rho(x)dx \qquad (6)$$

This expression clarifies the idea that the mean volume can change either by changing P(V|x) - e.g. changing the birth and division volume - or by changing ρ(x). Assuming ρ(x) is conserved, the only possibility which remains is to change P(V|x). This still does not *require* changing the birth/division volume. In particular, we examined whether the difference between a linear growth law, an exponential growth law:

$$P(V|x) = N\left(V, V_{Birth}\exp\left(\ln\left(\frac{V_{Division}}{V_{Birth}}\right)x\right), y\right),\ N(V, V_{Birth} + (V_{Division} - V_{Birth})x, y) \quad (7)$$

or a hypothetical logarithmic growth law:

$$P(V|x) = N\left(V, V_{Birth}\ln\left(\exp\left(\frac{V_{Division}}{V_{Birth}}\right)x\right), y\right) \quad (8),$$

where $N(V, x, y)$ indicates the normal distribution:

$$\mathrm{N}(V, x, y) = \frac{1}{\sqrt{2\pi(xy)^2}} \exp\left(-\frac{(V-x)^2}{2(xy)^2}\right) \qquad (9)$$

with a mean of $x$ and a coefficient of variation (CV) of $y$ could result in measurably different volume distributions. We examined the linear and exponential cases in the main text (Fig. 3c) and found the differences in the predicted volume distributions to be negligible. Here we provide an additional investigation of all three cases. Differing cell cycle durations do not impact the final volume distributions (the age distributions for the examples shown in the main text are included in Fig. S3a for completeness) and will be ignored.

Roughly doubling over the course of the cell cycle, we consider cells obeying logarithmic (dotted line), exponential (dashed line), and linear (solid line) growth laws increasing in size from one to two (red curves) and two to four (black curves) arbitrary units Fig. S3c. The resulting volume distributions depend additionally on the choice of CV – the larger the CV the greater the heterogeneity of the population. For CV larger than 0.15 which replicate the shape of the experimentally observed volume distributions and are displayed in the main text, the choice of growth law only modestly impacts the final volume distributions. See Fig. S3d. For smaller CV, however, the effects are more dramatic. See Fig S3e. In this small CV regime, the logarithmic growth law produces a significantly right-weighted volume distribution with cell spending a greater portion of the cell cycle at a larger volume. Similarly, the exponential growth law is left skewed and the linear growth law takes on the shape of the cell cycle completion distribution (Fig. S3b).

We may further utilize this framework to arrive at an approximation for the average ensemble volume in terms of the cell cycle duration, τ, and the volumetric growth rate $\frac{dV}{dt}$ alone. The result is similar for both exponential and linear cases so we will assume a linear growth law for simplicity. Now we may consider that a cell roughly doubles over the course of the cell cycle implying:

$$\mathrm{V_{Division}} = V_{Birth} + \tau\frac{dV}{dt} = 2\mathrm{V_{Birth}} \qquad (10)$$

and thus:

$$\overline{\mathrm{V}} = \int_0^1 [\mathrm{V_{Birth}}(1+\mathrm{x})] 2\ln(2)\exp(-\ln(2)\mathrm{x})\, dx = \frac{1}{ln(2)} V_{Birth} = \frac{1}{ln(2)}\tau\frac{dV}{dt} \approx \frac{3}{2}\tau\frac{dV}{dt} \qquad (11)$$

---

**3D cell shape reconstruction**

We sought to determine if increasing YAP/TAZ activity impacted cell shape. From previous work (Perez-Gonzalez et al., 2018), we know that cell geometry plays a major role in the force balance at the cell surface and therefore the cell shape. First traditional 2D projected shape was compared across the cell lines investigated with little variation observed due to the relatively non-protrusive behavior of the parental cell line, HEK293A. Next, we examined 3D shape where the apical surface was reconstructed from the epifluorescent images used to calculate volume. To collapse self-scaling (similar) cell shapes, the surfaces were normalized by cell volume (x, y, and z position were all scaled by the cubed root of the volume) so that each normalized surface enclosed a volume of 1 arbitrary unit. Being largely symmetrical, the cell lines investigated were found to be well approximated by spherical caps. The volume for a cap is given by the expression:

$$V = \frac{\pi r^3 \beta^2}{3}(3 - \beta) \qquad (12)$$

where the height of the cap is $r\beta$. The height distribution for each cell as a function of the distance from the center of mass of the cell (in this case center of volume) was calculated. A median height curve was then constructed and the best fit $\beta$,*r* were chosen for that curve. The 25th and 75th percentiles in $\beta$ were identified (representing the most "pancake-like" and the most hemispherical respectively). We found that there was no consistent trend in cell shape with increasing YAP/TAZ activity. The YAP/TAZ dKO, YAP KO, TAZ KO, and parental line (HEK293A) were all found to be self scaling (despite volume variation, the cell shapes were found to be similar when viewed as collective ensembles). It is interesting to note, however, the LATS1/2 dKO does not exhibit the same shape, with a significantly smaller mean $\beta$ (more "pancake-like", Fig. S3f**)**.

---

**G1/S Cell Size Checkpoint**

Another potential avenue through which changes in YAP/TAZ activity could be regulating cell volume is the specification of a G1/S cell size checkpoint. We looked at the birth volume, division volume, growth rate, DNA distribution, and pMLC data collected to determine if this possibility was self-consistent with previous measurements. We have observed the DNA distributions to be unchanged across the cell lines investigated, indicating the fraction of the cell cycle spent in G1 is likely to be conserved $t_{G1}/t_{Cycle} \equiv C$. Additionally, we found no significant dependence on cell cycle phase (e.g. G1, S, G2) for the volume growth rate indicating the growth curve may be well approximated (within one doubling) with a linear function $V = V_{Birth} + \frac{V_{Division} - V_{Birth}}{t_{Cycle}} t$. Thus, we may expect $\frac{V_{G1} - V_B}{V_D - V_B} \equiv C$. Noting that the cell roughly doubles in volume over the course of the cell cycle this may be simplified to be $\frac{V_{G1}}{V_B} \equiv C'$. We then found the lowest value of pMLC below which 90% of the G1 cells in each population lie, $PMLC_{G1}$. We found mean $V_B$ to increase roughly 30% from the smallest population, YAP/TAZ dKO, to the largest population, LATS1/2 dKO; however, $PMLC_{G1}$ increased two-fold (Fig. S5a). This implies the correspondence between pMLC and volume cannot be simply $V = \alpha pMLC$ but rather $V = \gamma + \alpha pMLC$ with a positive $\gamma$. We then found the best fit line for the mean volume across each population as a function of the mean pMLC and confirmed a positive $\gamma$ (Fig. 5e). Thus, all data collected is consistent with the possibility of YAP/TAZ-dependent scaling of G1 checkpoint volumes across the cell lines examined.

We went on to probe the effect of cell tension on the G1 checkpoint volume through the use of ROCK-inhibitor Y-27632**.** We found the progression from G1 to S was delayed with the addition of Y-27632. These results may be interpreted to indicate a lengthening of the G1 duration. Seeking to compare the experimental DNA distributions obtained with those expected through a predicted lengthening of G1, we constructed the following theoretical framework.

At the beginning of the experiment, cells occupy some initial cell cycle completion distribution, $\rho(x)$ (where "x" will be denoted the cell cycle "completion-fraction") and cell cycle duration distribution, $w(a)$. See "Computing Cell Volume Distributions" above. The cell cycle completion distribution may be propagated forward in time given the conditional probability of arriving at a new completion-fraction given that a cell was at a preceding completion-fraction at the previous timepoint. Assuming the time elapsed is sufficiently short so that no cell divides twice, the probability of arriving at completion-fraction *y* given a start at completion-fraction *x* takes one of two forms depending on whether a cell divides during the timestep. For the case with no division occurring during the specified timestep, this expression takes the form:

$$P(y(t)|x)dy = w\left(\tau = \frac{\Delta t}{y-x}\right)d\tau \qquad (13)$$

The case with a single cell division occurring during the specified timestep:

$$P(y(t)|x)dy = \left[\int_0^\infty w\left(\tau = \frac{\Delta t - (1-x)\tau_1}{y}\right)w(\tau_1)d\tau_1\right]d\tau \qquad (14)$$

where $\tau_1$ is the duration of the completed cell cycle. The total conditional probability is not simply the sum of the two terms above because the second term accounts for events during which the cell divides and thus are duplicated in the resultant distribution. Thus, the resultant distribution, $\rho_{\Delta t}(y)$ which is the cell cycle completion distribution at time $\Delta t$, is given by the following expression:

$$\rho_{\Delta t}(y) = A\left\{\int_0^y w\left(\tau = \frac{\Delta t}{y-x}\right)\frac{d\tau}{dy}\rho(x)dx + 2\int_0^1\left[\int_0^\infty w\left(\tau = \frac{\Delta t-(1-x)\tau_1}{y}\right)w(\tau_1)d\tau_1\right]\frac{d\tau}{dy}\rho(x)dx\right\} \qquad (15)$$

where $\rho(x)$ is the cell cycle completion distribution at time zero and the constant *A* indicates this expression requires renormalization. While this result may be achieved numerically, artifacts and instability largely due to the inverse relationship between y and $\tau$ make this cumbersome. Alternatively, the following approximation may be implemented

$$\int_0^\infty w\left(\tau = \frac{\Delta t-(1-x)\tau_1}{y}\right)w(\tau_1)d\tau_1 \sim w\left(\tau = \frac{\Delta t}{(1+y)-x}\right) \qquad (16)$$

While this is true for the case where $w(\tau) = \delta(\tau - \tau^*)$ it is not true in general; however, it is adequate for the purpose of this work and the following expression for the resulting distribution:

$$\rho_{\Delta t}(y) = A\left\{\int_0^y w\left(\tau = \frac{\Delta t}{y-x}\right)\frac{d\tau}{dy}\rho(x)dx + 2\int_0^1 w\left(\tau = \frac{\Delta t}{(1+y)-x}\right)\frac{d\tau}{dy}\rho(x)dx\right\} \qquad (17)$$

utilizing this approximate may be used.

While this expression may be used to propagate the cell cycle completion distribution forward in time, to make predictions about the distribution of cells residing in the cell cycle phases, $G1\ \ , S, and\ G2$ , we need to specify the completion-fraction at which each phase begins and ends. In principle, these may not be fixed quantities. For example, if a cell spends much more than the average amount of time in $G2$ in a given cycle, its daughter cells may spend less time in G1 to compensate. The model below does not include any potential mother-daughter correlations and simply assumes $\mathrm{G1}, S, and$ G2 occupy constant fractions of the cell cycle across the population until acted upon by the drug, $\mathrm{G1}: x \in [0, g_1]$; $S: x \in [g_1, s]$; $\mathrm{G2}: x \in [s, 1]$. The action of the drug is also simplified to be discrete. Before the drug is introduced, $\mathrm{G1}, S, and$ G2 are assumed to occupy constant fractions $g_1$, $s - g_1$, and $1 - s$. of the cell cycle. The moment the drug is introduced, these fractions are assumed to become $\frac{(1+\phi)g_1}{1+\phi g_1}$, $\frac{s-g_1}{1+\phi g_1}$, and $\frac{1-s}{1+\phi g_1}$ modelling a process by which the duration of G1 increases by a constant fraction $\phi$ and the durations of $S, and$ G2 are unchanged, $\mathrm{G1}: x \in [0, (1+\phi)g_1]$; $S: x \in [(1+\phi)g_1, s + \phi g_1]$; $\mathrm{G2}: x \in [s + \phi g_1, 1 + \phi g_1]$.

The cell cycle completion distribution is thus modified after the action of the drug to become $\rho_\phi(x)$. The map between $\rho(x)$ and $\rho_\phi(x)$ is assumed to be a uniform dilation of the interval $[0, g_1]$ to $[0, (1+\phi)g_1]$ and a rigid translation of the intervals $[g_1, s]$ and $[s, 1]$ to the intervals $[(1+\phi)g_1, s + \phi g_1]$ and $[s + \phi g_1, 1 + \phi g_1]$ respectively. In other words, cells in G1 experience a uniform dilation of the remaining portion of $G1$ they have yet to complete while cells in $S, and$ G2 are unaffected until the next division. These assumptions yield the following: when $x < (1+\phi)g_1$, $A\rho_\phi(x) = \rho\left(\frac{x}{1+\phi g_1}\right)$, when $x \geq (1+\phi)g_1$, $A\rho_\phi(x) = \rho(x - \phi g_1)$ and similarly, $A w_\phi(\tau) = w\left(\frac{\tau}{1+\phi g_1}\right)$ where $A$ indicates these expressions must be renormalized. Thus, the cell cycle completion distribution may be propagated forward during a timestep in which the drug acts using the following expression:

$$\rho_{\Delta t}(y) = A\left\{\int_0^y w_\phi\left(\tau = \frac{\Delta t}{y - x}\right)\frac{d\tau}{dy}\rho_\phi(x)dx + 2\int_0^1 w\left(\tau = \frac{\Delta t}{\left((1+\phi g_1) + y\right) - x}\right)\frac{d\tau}{dy}\rho_\phi(x)dx\right\} \quad (18)$$

---

**Cell Size Assessments.** Cells were sized and counted with a Coulter Counter (Multisizer 3, Beckman Coulter, Fullerton, CA) using an orifice size of 50 μm and a lower size measurement limit of 1 μm. In addition to the Coulter Counter measurements, cells were alternatively used by flow cytometry with the following protocol: First, cells were resuspended in DMEM with 10% FBS and 1% P/S. The sample was centrifuged at 1500 rpm for 5 minutes. The supernatant was removed, and the cells were resuspended in 5 mL of PBS into the tube. After this, cells were counted using a hemocytometer. We centrifuged 1 mL with 500,000 cells at 1500 rpm for 5 minutes. We proceeded to remove the supernatant and resuspended cells into 500 μL of ice-cold PBS. Afterwards, we added 4.5 mL of ice-cold 70% ethanol in 0.5 mL increments and vortexed in every iteration. This was followed by placing the cells in ice or freezer overnight. After this, we centrifuged again at 1500 rpm for 5 minutes to later remove the supernatant and resuspend cells in 1 mL PBS. To have nuclear signal we pipetted 10 μL of Stock Hoechst 33342 (final Hoechst concentration should be 100 μg/mL) and incubated cells for 60 minutes. Finally, we transferred the mix into a test tube with a Corning Falcon Test Tube with Cell Strainer Snap Cap (i.e. filter) and measured the fluorescence intensity using a SH800S Cell Sorter.

**Micro-fluidic device fabrication.** Silicon molds were fabricated using standard photolithography procedures. Masks were designed using AutoCAD and ordered from FineLineImaging. Molds where made by following manufacturer's instruction for SU8-3000 photoresist. Two layers of photoresist were spin coated on a silicon wafer (IWS) at 500 rpm for 7 seconds with acceleration of 100 rpm/s and 2000 rpm for 30 seconds with acceleration of 300 rpm/s respectively. After a soft bake of 4 minutes at 95 $^{o}$C UV light was used to etch the desired patterns from negative photoresist to yield feature heights that were approximately 15 μm. The length of the abovementioned channels is 16.88 mm and the width is 1.46 mm.

A 10:1 ratio of PDMS Sylgard 184 silicone elastomer and curing agent were vigorously stirred, vacuum degassed, poured onto each silicon wafer and cured in an oven at 80 $^{o}$C for 45 minutes. Razor blades were then used to cut the devices into the proper dimensions, inlet and outlet ports were punched using a blunt-tipped 21 Gauge needle (McMaster Carr, 76165A679). The devices were then sonicated in 100% IPA for 15 min, rinsed with water and dried using a compressed air gun.

50 mm glass bottom petri-dishes (FlouroDish Cell Culture Dish, World Precision Instruments) were cleaned with water and then dried using a compressed air gun. The petri-dishes and PMDS devices were then exposed to oxygen plasma for 1 minute for bonding. Finally, the bonded devices were placed in an oven at 80 $^{o}$C for 45 minutes to further ensure enhance bonding.

**Cell Volume Measurements.** Micro-fluidic chambers were exposed to 30s oxygen plasma before being incubated with 50 μg/mL of type I rat-tail collagen (Corning; 354236) for 1 hour at 37 $^{o}$C. The chambers were washed with 1X PBS before approximately 50,000 cells were injected into them. The dishes were then immersed in media to prevent evaporation. The cells were seeded with

0.1 µg/mL of Alexa Fluor 488 Dextran (MW 2000 kD; ThermoFisher), allowed to adhere in the incubator at 37 °C with 5% $CO_2$ and 90% relative humidity and then imaged within 12 hours.

The cells were imaged using a Zeiss Axio Observer inverted, wide-field microscope using a 20x air, 0.8 numerical aperture (NA) objective equipped with an Axiocam 560 mono charged-coupled device (CCD) camera. The microscope was equipped with a $CO_2$ Module S (Zeiss) and TempModule S (Zeiss) stage-top incubator (Pecon) that was set to 37 °C with 5% $CO_2$ for long-time imaging. Differential interference contrast (DIC) microscopy was used to accurately capture the cell area and shape and Epifluorescent microscopy was used to measure volume. Individual cells were traced using the following algorithm.

Cell contours were segmented from the DIC and epifluorescence (Volume) channels. First, a rough contour is generated from a smoothed copy of the Epi channel where pixels darker than the background intensity are identified. Next a measure of the local contrast of the DIC channel (here high contrast regions are identified) is used to expand the contour to include small features (small lamellipodia etc.) which have low contrast in the Volume channel and may be missed. This expanded contour is used to identify the cell boundary. Inner and outer annuli are created by dilating this contour 10 and 25 pixels away from the cell (Inner boundary shown in figure S1a). The mean fluorescence intensity of the pixels between inner and outer annulus, or mean background intensity, $I_{annulus}$, is related to the total channel height. The volume boundary, shown as purple line in Figure S1b, is created by dilating the cell contour 20 pixels away from the cell. The local fluorescence intensity enclosed by the volume boundary, $I_V$, corresponds to the local height above the cell ($h_2$, shown in Figure 1(a), main text). The volume of the cell is then calculated as follows: $V = Channel\ Height\ \sum_{pixels\ within\ volume\ boundary} \left(1 - \frac{I_V}{I_{Annulus}}\right) \delta A$.

Every experiment on cell volume was repeated at least three times with three technical repeats corresponding to the three individual channels in the microdevice. Experiments in glass gave at least 50 single cell measurements. Softer substrates yielded smaller datasets per measurement. The sample size for volume measurements was kept over 200 single cells with the exception of birth and peak volume on fig. 1n. This was done in order to get a normal distribution for each complete dataset.

**Immunofluorescence.** Immunofluorescence was carried out as described as in (Aifuwa et al, 2015). Briefly, cells were seeded at either single cell density (12,000 cells/cm$^2$ for all HEK 293A cell lines) for 6 hrs and then fixed with 4% paraformaldehyde (100503-917, VWR) for 10 minutes. Samples were then rinsed 3 times with 1X PBS. 0.1% Triton X-100 (T8787, Sigma Aldrich) dissolved in PBS is then added for 10 minutes, washed 3 times with 1X PBS and then the fixed cells are blocked with 1% bovine serum albumin (A7906, Sigma Aldrich) for 1 hour at room temperature. Primary antibodies are incubated overnight in 1% BSA. Antibodies used included: YAP 63.7 (1:100; ms; SC/101199), Phospho-Myosin Light Chain 2 Thr18/Ser19 (1:100; rb; Cell Signaling Technology #3674), pS6 (1:1000, rb, Cell Signaling Technology #5364). The next day the dishes are rinsed 3 times with 1X PBS and incubated for 2 hrs in secondary antibodies with the following secondary antibodies Mouse Alexa Fluor 488, Rabbit 568, and DNA was stained using 20 μg/mL of Hoechst 33342. In addition, in combination with the Hoechst 33342, we used a succinimidyl ester dye (SE-A647), which reacts with lysil groups reported by Kafri et al. (2013).

Wide-field microscopy using the set-up described above was used to measure the total pMLC, YAP/TAZ, pS6, DNA content, and total protein content of the cells. To obtain spatial information about pMLC we used a Zeiss LSM 800 confocal microscope equipped with a 63X oil-immersion, 1.2 (NA) objective. A 567nm laser was used to image the stained cells. Images were acquired with a resolution of 1024 x 1024, which gives a field of view of 10485.76 μm$^2$. We imaged the cells with confocal image stocks of total thickness of 20 μm to cover the entire height of the cells. Confocal image slices were spaced 2 μm apart and the pinhole size was 1 μm.

For each fluorescence image, we subtract the pixel intensities with mean background intensity. A binary mask is generated based on the pixel intensities of fluorescence image (for the pixel intensities within the cell region is much higher than the intensities of anywhere else), where pixels within the cell/nucleus region are marked with “1” and pixels outside the cell/nucleus are marked with “0”. By multiplying the binary mask with actual florescence image, we can identify all the pixel values that is within the cell/nucleus. The total intensities within cell/nucleus boundary is calculated by summing up all the intensity values. The cell and nucleus boundary is then traced by Matlab routine “bwboundaries”. Every traced region with total area of 1,500 pixels square or less is considered as debris or cell fragments, and, therefore, is ignored.

We utilized the pMLC channel to generate the binary mask for the cell. The traced boundary is then dilated 15 pixels away from the cell, to capture all the scattered light from epifluorescence image. The binary mask for the cell nucleus is generated based on Hoechst channel. No dilation is made on nucleus mask, to avoid overestimation of total nucleus YAP. We multiply the nucleus mask with every cell mask, to exclude all the nuclei from other cells within the same field of view. The traced boundary is shown in Fig. S1 (b).

For confocal *z* stacks, the basal layer of the cell is identified when clear stress fibers are seen (as example shown in Fig. S5 (b)). All stacks that are below the basal layer are neglected. We identified the first apical slide when the stress fibers disappear. The traced boundary of every apical slides is dilated 5 pixels (~ 1 micrometer) inside the cell, to mark the inner boundary of cortical layer. Fig. 5d shows pMLC are mainly cortical, except for basal layer, where some stress fibers can be seen. Therefore, the pMLC within the cell cytoplasm is very minimal compared to cortical pMLC.

Every experiment was repeated two times with two technical repeats on every experiment. In addition, each technical repeat consisted of at least 100 single cell measurements (except for confocal measurements). The sample size for qIF aimed for at least 200 single cells. This dataset size was targeted in order to get a normal distribution for each complete dataset. Finally, no single cells were excluded during the analysis of these datasets. The only cells excluded we those forming clusters.

**Cell Protein Synthesis Measurement**. SUnSET method (Schmidt et al, 2009) was applied as a measurement of single cell protein synthesis rate. The HEKs were seeded 20,000 cells/mL in a 24-well plate for 4 hours in the incubator with DMEM (10% FBS, 1% PS) media, and then treated with 10 µg/mL puromycin (P8833, Sigma Aldrich) diluted in dPBS for 10 minutes in the incubator. Cells were fixed using 4% PFA right after puromycin treatment and stained according to immune-fluorescence protocol described above. Anti-puromycin antibody, clone 12D10 (MABE343 EMD, Millipore) was used in the ratio of 1:1000 in BSA as the primary antibody solution and Mouse Alexa Fluor 488 was used in the ratio of 1:1000 in dPBS as the secondary antibody solution.

**Western Blotting.** Cells were lysed in SDS sample buffer (50mM Tris pH 6.8, 2% SDS, 0.025% bromophenol blue, 10% glycerol, 5% Beta mercaptoethanol), and boiled for 5 minutes. Proteins were separated on 8% to 10% Bis-Tris polyacrylamide gels. Immunoblots were performed as previously described (Meng et al., 2015). Antibodies for Lats1 (#9153) and Lats2 were purchased from Cell Signaling Technology Lats2 (5888). YAP 63.7 (sc-101199) was purchased from Santa Cruz (this antibody recognizes both YAP and TAZ). Vinculin (V9131 was purchased from BD Biosciences).

**Statistical Analysis.** C - To show significance we used a one way non-parametric anova (we did not assume gaussian distributions given the shapes of the histograms). We performed the Kruskal-Wallis test. We also performed follow up tests comparing the mean rank of each column with the mean rank of a control column of HEK293A. We performed Dunn's multiple comparisons test and obtained the corresponding P-value.

For comparison between two groups such as the birth volume and peak volume in figure 1n, we performed an unpaired non-parametric Mann-Whitney test. For comparison between protein-expression experiments via qIF, we used a one-way anova analysis with a Brown-Forsythe test.

**SM Figure Captions**

**Figure S1.** (a) DIC image overlaid with cell boundary used for calculating morphological properties. (b) Epifluorescent volume image overlaid with the boundary used to segment the region integrated to yield volume as described in the main text. The volume boundary is dilated 20 pixels out from the cell boundary. The annulus used to calculate the background intensity is constructed from boundaries dilated 10 and 25 pixels from the cell boundary (not shown). (c) Three-dimensional reconstruction of the cell. The heights are derived from the epifluorescent volume image as described in the main text and colorized with the intensity map defined by the DIC image. (d) DIC image of a fixed cell. (e) Immunofluorescent image of Hoechst labeling with nuclear boundary overlaid. (f) Immunofluorescent image of YAP/TAZ labeling with both the nuclear boundary and cell boundary overlaid. Scale bars are 20 μm. (g) Volume versus area for each one of the five cell lines including the CRISPR knockouts. ($N_{YAPTAZ\ dKO}$=70, $N_{YAP\ KO}$=81, $N_{TAZ\ KO}$=95, $N_{HEK\ 293A}$=74, $N_{LATS1/2\ dKO}$=61). (h) shows the distribution for cell area for each cell line from the previous figure S1g.

**Figure S2.** (a) Growth rate characterizing volume increase at single cell level by using a linear fitting to each individual curve. The first five panels show the individual cell growth versus their associated initial volume. The last panel shows the slope of the fitting for all five cell lines ($N_{YAPTAZ\ dKO}$=130, $N_{YAP\ KO}$=118, $N_{TAZ\ KO}$=162, $N_{HEK\ 293A}$=175, $N_{LATS1/2\ dKO}$=185). (b) Growth rate characterizing volume increase at single cell level with an alternative representation. The five panels show the individual cell growth versus their volume over time ($N_{YAPTAZ\ dKO}$=130, $N_{YAP\ KO}$=118, $N_{TAZ\ KO}$=162, $N_{HEK\ 293A}$=175, $N_{LATS1/2\ dKO}$=185).

**Figure S3.** (a) Example cell age distributions for cells with a long cell cycle duration (red line) and a short cell cycle duration (black line). (b) The cell cycle completion distribution well conserved for a variety of cell cycle distributions. (c) Growth trajectories for cells obeying a hypothetical logarithmic growth law (dotted lines), linear growth law (solid line), and exponential growth law (dashed line). Small cells ranging in volume from one to two arbitrary units (red curves) and large cells ranging in volume from two to four arbitrary units (black curves) are displayed. (d) Resultant volume distributions where the conditional probability has a large CV. All growth laws produce similar volume distributions. (e) Resultant volume distributions where the conditional probability has a small CV. The resulting volume distributions are clearly distinguishable. (f) The fitted spherical cap distributions for the Hippo knockouts (red) in comparison to control HEK 293A (gray). The shaded band represents the median 50% of the population. The only cell line with a significant shape difference is the LATS1/2 dKO.

**Figure S4.** (a) IC50 curve for rapamycin exposure on the parental cell line. mTOR activity as measured by total pS6 decreases as the concentration of rapamycin is increase ($N_{Control}$ =111, $N_{10\ pM}$ =150, $N_{100\ pM}$=198, $N_{500\ pM}$=116, $N_{1\ nM}$ =178). (b) IC50 curve for rapamycin exposure on the parental line. YAP/TAZ expression remains roughly constant through exposure to rapamycin. (c) Effect of rapamycin on cell volume at 4 hours, 24 hours, 72 hours and 216 hours. Left shows average statistics while left shows all data. ($N_{Control}$ =314, $N_{Rapamycin\ 4\ hrs}$ =224, $N_{Rapamycin\ 24\ hrs}$ =46, $N_{Rapamycin\ 72\ hrs}$ =289, $N_{Rapamycin\ 216\ hrs}$ =28). (d) Effect on mTOR activity by rapamycin for all 5 cell lines at 4 hrs and 72 hrs. ($N_{YAP/TAZ\ dKO\ Control}$ =11, $N_{YAP/TAZ\ dKO\ Rapa\ 4hrs}$ =33, $N_{YAP/TAZ\ dKO\ Rapa\ 72\ hrs}$ =38, $N_{YAP\ KO\ Control}$ =117, $N_{YAP\ KO\ Rapa\ 4hrs}$ =36, $N_{YAP\ KO\ Rapa\ 72\ hrs}$ =85, $N_{TAZ\ KO\ Control}$ =90, $N_{TAZ\ KO\ Rapa\ 4hrs}$ =157, $N_{TAZ\ KO\ Rapa\ 72\ hrs}$ =93, $N_{Parental\ Control}$ =46, $N_{Parental\ Rapa\ 4hrs}$ =43, $N_{Parental\ Rapa\ 72\ hrs}$ =68, $N_{LATS1/2\ dKO\ Control}$ =7, $N_{LATS1/2\ dKO\ Rapa\ 4hrs}$ =92, $N_{LATS1/2\ dKO\ Rapa\ 72\ hrs}$ =90). (e) YAP/TAZ expression in the 5 cell lines with rapamycin treatment. (f) Average volume of the five cell lines in the Hippo pathway (red) and average volume of the five cell lines with rapamycin treatment (pink).

**Figure S5.** (a) Average pMLC expression across population of HEKs over 10 z-stacks. Each stack is 1 μm interval. The bottom stack (stack #1) is manually chosen and focused to qualitatively match the maximum radius of the adherent surface of a cell. (b) Apical and basal pMLC expression across HEKs. The top-left panel shows the apical pMLC expression for the single cells of HEKS. The top-right panel shows the basal pMLC expression for the single cells of HEKs. The bottom panel shows the Apical/Basal ratio of pMLC expression for the single cells of HEKs. The bottom 2 stacks are counted as the basal stacks and the top 8 stacks are counted as apical stacks. The sum of the fluorescent signal from corresponding stacks are used to report the expression of apical or basal pMLC. ($N_{YAPTAZ\ dKO}$=40, $N_{YAP\ KO}$=36, $N_{TAZ\ KO}$=36, $N_{HEK\ 293A}$=37, $N_{LATS1/2\ dKO}$=43). (c) Example of the pMLC signal over different z-stacks. A HEK-293A parental cell was investigated and the pMLC confocal fluorescence images over 6 z-stacks from the bottom of this cell were shown.

### Figure S1

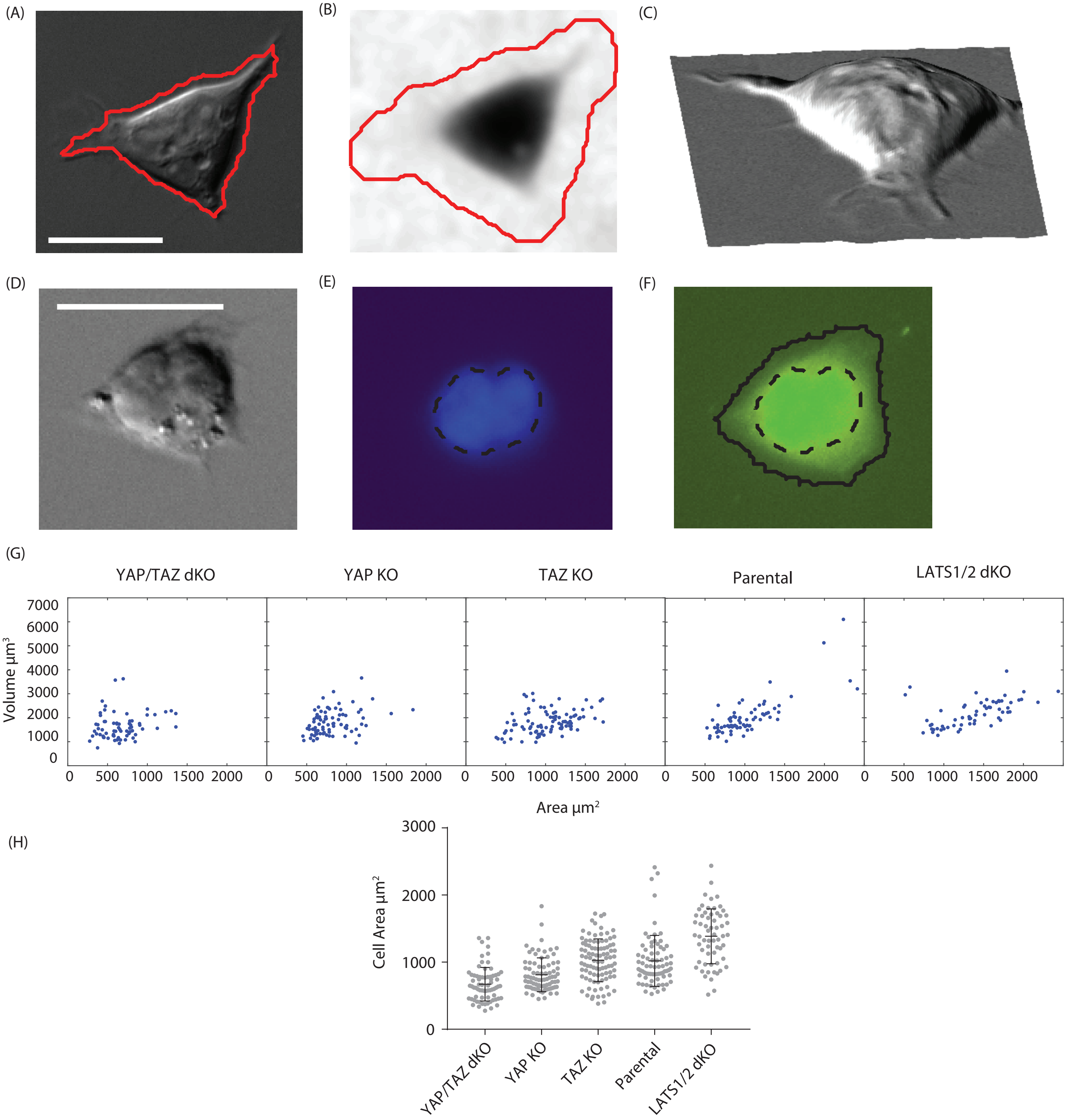


| | YAP/TAZ dKO | YAP KO | TAZ KO | Parental | LATS1/2 dKO |
|---|---|---|---|---|---|
| Average Area ± error | 671.6 ± 29.9µm² | 811.3 ± 27.4 µm² | 1028 ± 32.7 µm² | 1018 ± 44.2 µm² | 1386 ± 52.4 µm² |

Table 1. Statistics of Cell Area across the Hippo Pathway knockouts. Average Value + SEM.

| | YAP/TAZ dKO | YAP KO | TAZ KO | Parental | LATS1/2 dKO |
|---|---|---|---|---|---|
| Average Volume ± error | 1725 ± 38.4µm³ | 1931 ± 51.5 µm³ | 1986 ± 52.2 µm³ | 2362 ± 60.9 µm³ | 2437 ± 53.5 µm³ |

Table 2. Statistics of Cell Volume across the Hippo Pathway knockouts. Average Value + SEM.

Figure S2

(A) Linear fit

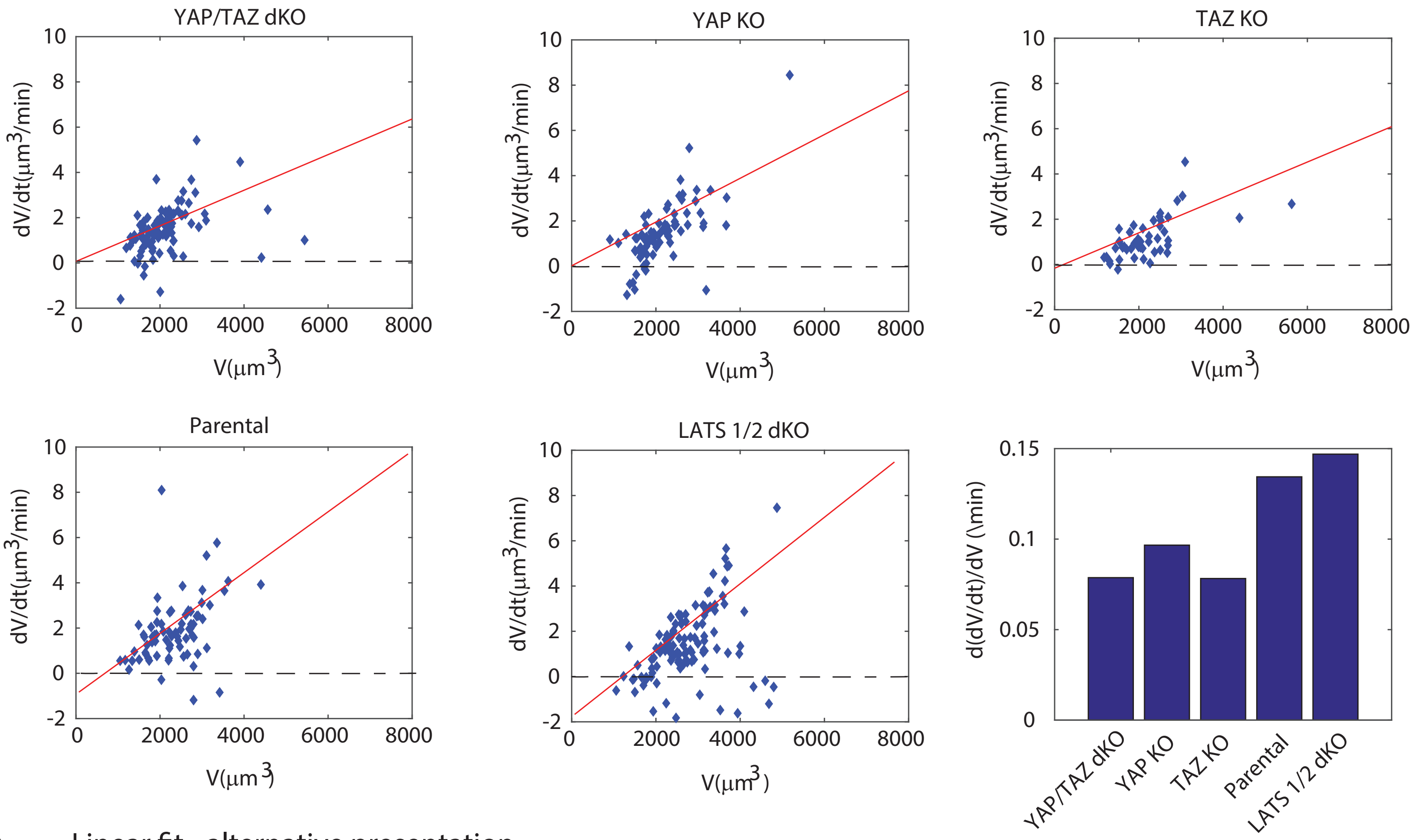


(B) Linear fit - alternative presentation

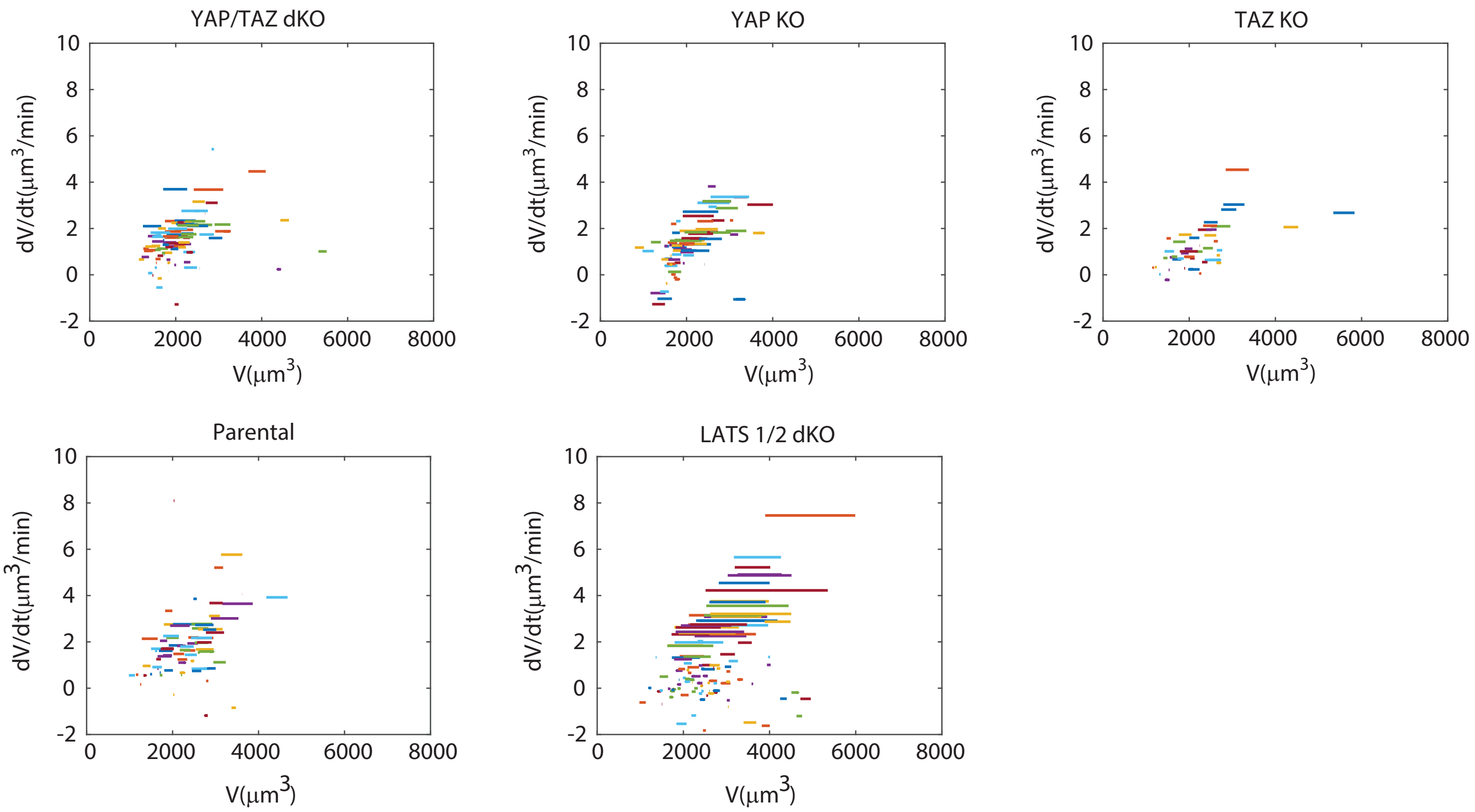

### Figure S3

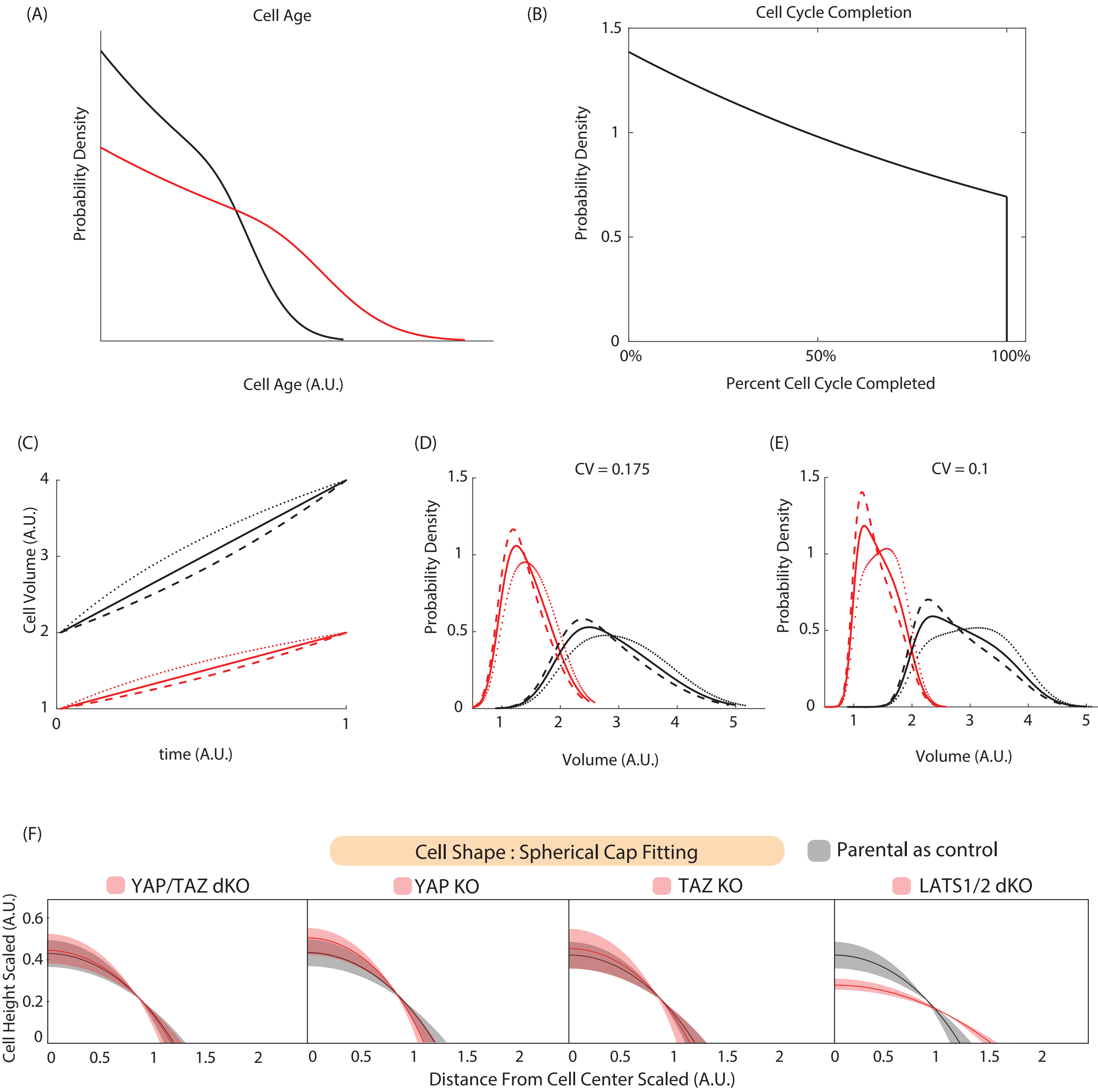

(A)
Cell Age
Probability Density
Cell Age (A.U.)
(B)
Cell Cycle Completion
1.5
1
0.5
0
0%
50%
100%
Probability Density
Percent Cell Cycle Completed
(C)
4
3
2
1
0
1
Cell Volume (A.U.)
time (A.U.)
(D)
CV = 0.175
1.5
1
0.5
0
1
2
3
4
5
Probability Density
Volume (A.U.)
(E)
CV = 0.1
1.5
1
0.5
0
1
2
3
4
5
Probability Density
Volume (A.U.)
(F)
Cell Shape : Spherical Cap Fitting
Parental as control
YAP/TAZ dKO
YAP KO
TAZ KO
LATS1/2 dKO
0.6
0.4
0.2
0
Cell Height Scaled (A.U.)
0
0.5
1
1.5
2
Distance From Cell Center Scaled (A.U.)

### Figure S4

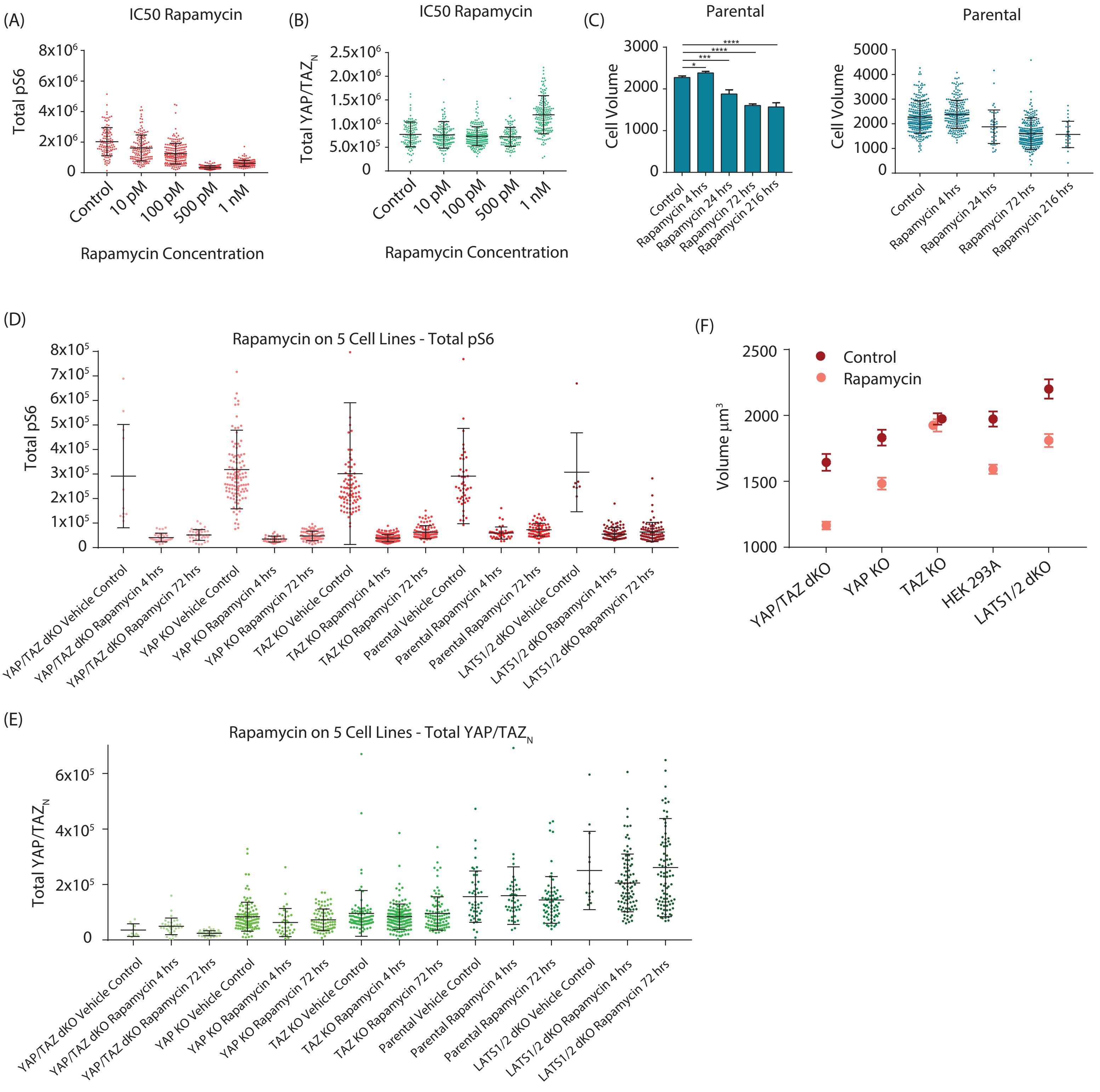


| | YAP/TAZ | YAP | TAZ | HEK 293A | LATS1/2 |
|---|---|---|---|---|---|
| Vehicle Control Average Volume (DMSO) | 1644 µm³ | 1832 µm³ | 1974 µm³ | 1973 µm³ | 2201 µm³ |
| Vehicle Control Standard deviation (DMSO) | 64 µm³ | 60 µm³ | 43 µm³ | 57 µm³ | 73 µm³ |
| 1nM Rapamycin Average Volume | 1165 µm³ | 1483 µm³ | 1925 µm³ | 1592 µm³ | 1810 µm³ |
| 1nM Rapamycin Standard deviation | 29.96 µm³ | 45 µm³ | 47.25 µm³ | 35.29 µm³ | 49 µm³ |

Table 3. Statistics of Cell Volume across the Hippo Pathway knockouts and its respective treatment with Rapamycin.

### Figure S5

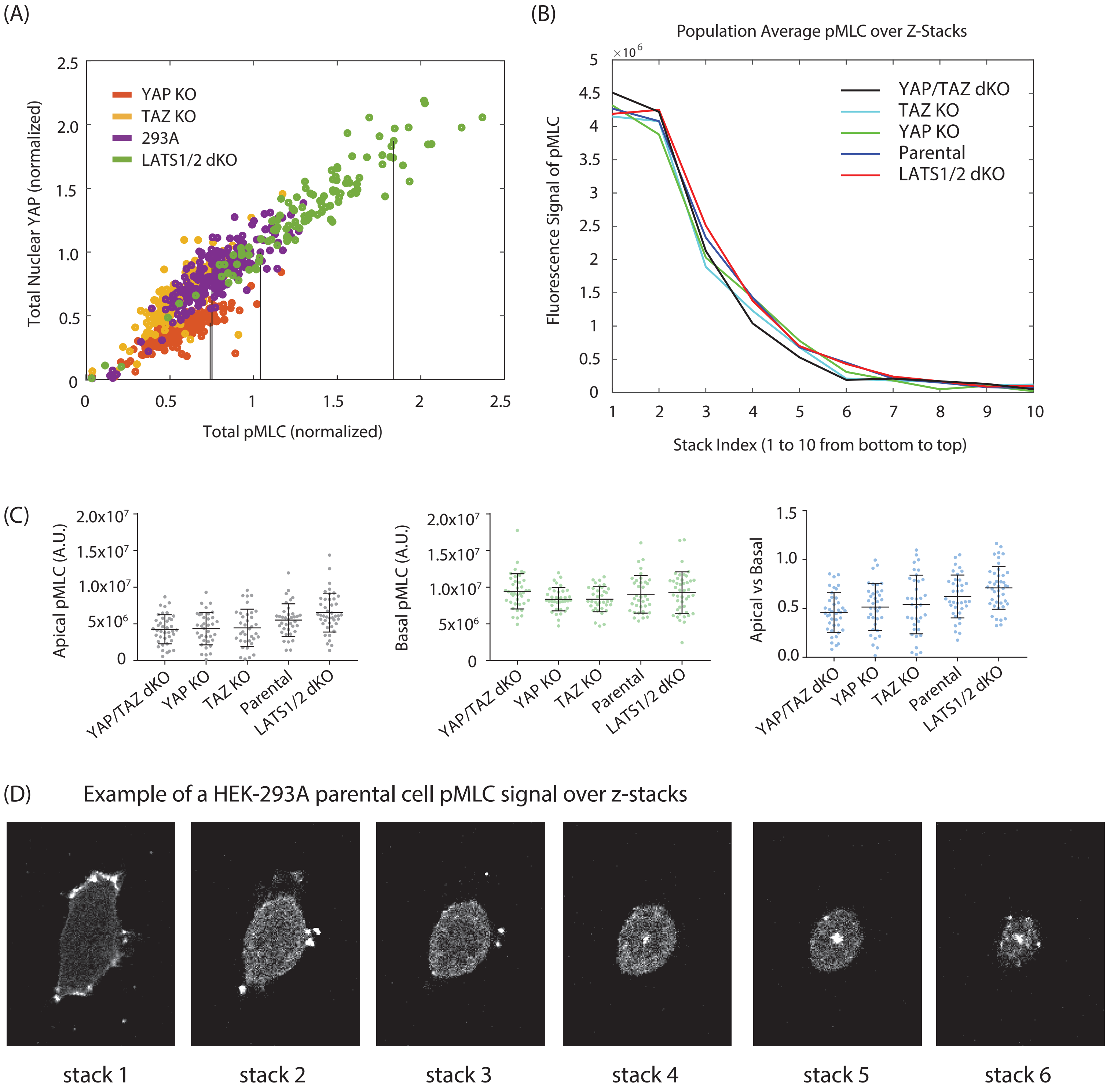

(A)
YAP KO
TAZ KO
293A
LATS1/2 dKO
Total Nuclear YAP (normalized)
Total pMLC (normalized)
0
0.5
1.0
1.5
2.0
2.5
0
0.5
1
1.5
2
2.5
(B)
Population Average pMLC over Z-Stacks
×10^6
Fluorescence Signal of pMLC
YAP/TAZ dKO
TAZ KO
YAP KO
Parental
LATS1/2 dKO
Stack Index (1 to 10 from bottom to top)
(C)
Apical pMLC (A.U.)
Basal pMLC (A.U.)
Apical vs Basal
2.0x10^7
1.5x10^7
1.0x10^7
5x10^6
YAP/TAZ dKO
YAP KO
TAZ KO
Parental
LATS1/2 dKO
(D)
Example of a HEK-293A parental cell pMLC signal over z-stacks
stack 1
stack 2
stack 3
stack 4
stack 5
stack 6
